# Mining conserved and divergent signals in 5’ splicing site sequences across fungi, metazoa and plants

**DOI:** 10.1101/2021.07.14.452117

**Authors:** Maximiliano Beckel, Bruno Kaufman, Marcelo Yanovsky, Ariel Chernomoretz

## Abstract

The main steps of the splicing process are similar across eukaryotes. However differences in splicing factors, gene architecture and sequence divergences suggest clade-specific features of splicing and its regulation. In each organism the ensemble of 5’ splicing sequences reflects the balance between natural nucleotidic variability and minimal molecular constraints to assure splicing fidelity. This compromise shapes the underlying statistical patterns in donor sequences composition.

In this work we aimed to mine conserved and divergent signals in splicing donor sequences. As 5’ donor sequences are a major cue for proper recognition of splicing sites we reasoned that statistical regularities of their sequence composition might reflect biological functionality and evolutionary history associated to splicing mechanisms.

We considered a regularized maximum entropy modeling framework to mine for non-trivial two-site correlations in donor sequences of 30 different eukaryote organisms. Our approach allowed us to accommodate and extend within a unified framework many of the regularities observed in previous works, like the negative epistatic effects between exonic and intronic consensus sites. In addition, for each analyzed organism, we could identify minimal sets of two-site coupling patterns that could generate, at a given regularization level, observed one-site and two-site frequencies in donor sequences. Noticeably, performing a systematic and comparative analysis of 5’ss we showed that lineage information could be traced from joint di-nucleotide probabilities. Specifically, we could identify characteristic two-site coupling patterns for plants and for animals and argue that they could echo differences in splicing regulation previously reported between these groups.

**Author summary:** The sequence composition of 5’ splicing sites of eukaryote organisms reflect a complex scenario. Nucleotide variability has to coexist with the need to correctly define exon/intron boundaries and the fidelity of splicing seems to depend on a pattern of trade-offs between substitutions at different positions of the splicing site. Better understanding of these patterns may help to gain insight into the details underlying the splicing process and its evolution.

In this work we aimed to study conserved and divergent signatures embedded in the sequence composition of eukaryotic 5’ splicing sites. We developed generative probabilistic models that allowed us to analyze sequence composition of donor sequences for several eukaryote organisms. Our regularized models served to incrementally disentangle the minimal set of coupling parameters needed to accurately reproduced observed 1-site and 2-site nucleotide frequencies. Focusing our study on di-nucelotide probabilities we found that they actually carried phylogenetic signal. In particular, our comparative analysis allowed us to identify differential two-site coupling patterns for animal and plants that might be related to specific differences in splicing regulation.

## Introduction

In most eukaryotic genes coding DNA regions - called exons - are interrupted by non coding ones. These sequences, named introns, are co-transcriptionally spliced-out from the nascent transcript in a process called splicing [1–3]. RNA splicing results from a coordinated and sequential set of biochemical reactions that involve small nuclear ribonucleoproteins (snRNPs) which, together with less stably associated non-snRNP proteins, conform a dynamical molecular machinery called spliceosome [1, 4]. Two type of spliceosomes are known to operate in eukaryotes. A major U2 type (with U1, U2, U4, U5, and U6 snRNPs) that processes the majority of pre-mRNAs and a minor U12 type (with U11, U12, U4atac, U5, and U6atac snRNPs) that splices a minor fraction of pre-mRNAs with so-called U12 type introns [5].

Despite some lineage-specific deviations, four major sequence cues serve to place the spliceosome at the right locations on the immature transcripts. For the vast majority of U2 spliceosomal introns, conserved GT and AG di-nucleotides are recognized at the beginning of an intron (5’ splice site or donor site) and at the opposite intronic end (3’ splice site or acceptor site) respectively. In addition, a branching point (BP), presenting a conserved A residue, is located 18 to 40 nucleotides upstream of the 3’ss. Finally, a poly-pyrimidine tract follows the BP and complete the necessary set of sequence cues used to guide the spliceosome assembly [5–7].

The 5’ splicing site is a major requirement for splicing to take place. These donor sequences are involved in a key step of RNA splicing reactions in which boundaries between exons and introns are recognized. At this step, U1 identifies the 5’ splice junction between adjacent exons and introns through a complex formation that depends on highly conserved base pairing between the 5’ splice site and the 5’ end of U1 snRNA. Specifically, U1 snRNA forms base pairs across intron-exon junctions, potentially base-pairing the last three positions of the exon (positions -3 to -1) and the first 6 positions of the intron (positions +1 to +6). This hybridization is stabilized by U1-C, one of the protein component of U1 snRNP, that sets up hydrogen bonds between its polypeptide chain and the sugar-phosphate backbone of the pre-mRNA. Being the action of U1-C sequence-agnostic it is particularly relevant to deal with splice site sequence variability [8]. Before the first splicing catalytic step, U1 is replaced by U5 and U6 snRNPs, for which the snRNA also needs to bind to the exonic (−3 to +1) and intronic (+5,+6) portions of the 5’ss, respectively [6, 9, 10]. Moreover, in a recent contribution Artemyeva & Porter reported that U5 might also play a significant role in exon-intron boundary recognition [11]. All these findings support the idea that, albeit gene contexts could be relevant, splicing efficiency depends of 5’ss recognition to a great degree [12].

Given the relevance of donor sites, their sequence composition has been the subject of many studies which aimed to uncover non-trivial sequence patterns associated to relevant biology. In particular, information theoretical approaches were widely used to shed light on biological functionality and/or to trace the evolutionary history of splicing. For instance, in the earlies 90, Stephens & Schneider used information theory to analyze 1800 human splice sites. They could quantitatively estimate position-dependent contributions using a Shannon-like information measure and found that more than 80% of the sequence information (i.e. sequence conservation) was confined in the intronic part of donors sites [13]. The position-dependent information content measure was also considered by Sverdlov and collaborators to uncover differences in the way the information was distributed between exonic and intronic parts of nucleotide sequences in new (i.e. lineage-specific) and old (shared by two or more major eukaryote lineages) introns. They reported that old introns presented lower information content in the exonic than the intronic part of donor sites and, conversely, the opposite trend was true for new introns. This observation suggested an evolutionary splice signal migration from exon to intron during evolution [14]. More recently, Iwata & Gotoh compared 61 eukaryote species, and shown that even though donor site motifs resembled each other (suggesting that the spliceosome machinery is well conserved among eukarya) they did exhibit some degree of specificity [15].

Single-site statistics, like logos or consensus sequences, do not exhaust the relevant statistics embedded in 5’splicing sequences. There have also been attempts to characterize higher correlation patterns in donor site sequences beyond one-site statistics. One of the first results of this kind was reported in the aforementioned Stephens & Schneider’s paper. Despite that no mechanistic interpretation was provided, using information theory they were able to find significant mutual information values, of around 0.05, 0.07 and 0.04 bits, between human 5’ss positions (−2,+4), (−1,+5) and (−2,+5) respectively [13]. Almost ten years later, and following a completely different route, Thanaraj & Robinson considered decision trees to predict exon boundaries and found that long range dinucleotide associations (−1,+5) and (−2,+5) carried significant splicing signals [16]. These observed couplings between (−1,+5) and (−2,+5) positions have also emerged in a human-mouse comparative genomic study carried on by Carmel and collaborators [17]. Two-point correlations in human 5’splice sites were also analyzed by Sahashi and collaborators. They reported that non-complementary nucleotides to U1 snRNA at specific positions were compensated by complementary nucleotides at other positions, suggesting that a stretch of complementary nucleotides either in an exonic or an intronic region was essential for proper splicing [18]. Denisov and collaborators presented evidence that supported and extended this idea. Through a comparative analysis of the genome of three mammals they found a well-defined pattern of epistatic interactions between the strength of nucleotides occupying different positions along the donor site. While the strength correlation within both, the intronic and the exonic part, were found to be positive (i.e. positive epistatsis), nucleotide strength correlations between intronic and exonic parts were found to present negative epistasis [19]. Another relevant work that studied RNA motifs in connection with splicing signals was presented by Yeo & Burge [20]. Considering an entropy maximization framework, they showed that maximum entropy distributions consistent with different sets of constraints (i.e. enforcing low-order marginal probability distributions to match empirical values) could be used to recognized true RNA splicing sites in primary transcript sequences. They proposed a likelihood ratio statistics to discriminate real splice sites from decoys and showed that relevant constraints could be identified studying the amount of entropy reduction they induced [20].

Models rooted in the principle of maximum entropy have been applied to several biological problems in the last decades [21, 22]. The success of this modeling strategy is based on the observation that low-order (mainly pairwise) correlations were found to play an important role in several biological systems, and accurate descriptions can be established from observed probability distributions over pairs of elements [23, 24]. In this contribution we wanted to delve further into donor sequence regularities following that route. Considering a maximum entropy framework we developed a generative probabilistic model that allowed us to analyze sequence composition of donor sequences for several eukaryote organisms. With this approach we aimed to look for lineage specific signatures in order to gain biological insights about splicing.

The paper was organized as follow. We first presented our modeling framework. We discussed how our regularized models accurately reproduced observed 1-site and 2-site nucleotide frequencies and served to incrementally disentangle the hierarchy of coupling parameters sufficient to reproduce observed correlations at a given precision. Then, several aspects related with the data-driven energy function naturally introduced in our maximum entropy approach were discussed. We focused on the characterization of the identified coupling patterns afterwards. A comparative analysis allowed us to identify robust and conserved signatures and extend previously detected epistatic signals in donor sites. We then showed that di-nucleotide two-site probabilities actually carried phylogenetic signal and identified particular coupling patterns that were specific for plants and for animals. We finally discussed some biologically relevant implications of our results and drew final conclusions in the last section of the manuscript.

## Materials and methods

### Analyzed genomes

We considered annotated genomes from 30 eukaryotic species, including 5 fungi, 8 plantae and 17 metazoan genomes (see Supplementary Table 1). A custom script was used to automatically extract 9-base length donor sequences based on provided genome sequences (FASTA files) and annotation (GTF/GFF3 files) downloaded from Ensembl.

### Statistical model

For each organism, we aimed to approximate the joint probability distribution function, 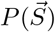, associated to the observed ensemble of 5′ splicing sites. Each element of this ensemble was a 9-base long sequence (last three exon’s positions followed by the first six intronic ones) 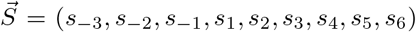 where *s*_*i*_ ∈ {*A, C, G, T*}. We worked under the hypothesis that the sought distribution should be compatible with observed 1-site and 2-site marginal probabilities, *f*_*i*_(*s*_*i*_), *f*_*ij*_(*s*_*i*_, *s*_*j*_), and implemented a maximization entropy approach to find the minimal structured distribution consistent with these constraints. Under this framework the estimated density probability function, 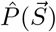, could be cast into a Boltzmann-like form (see [22, 25] for technical details):

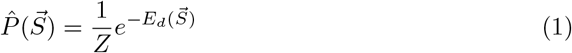

with

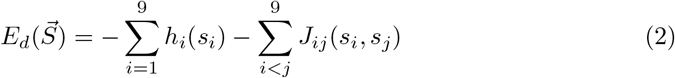

playing the role of a data-driven energy. *Z*, also known as the *partition function*, can be considered here a normalization constant. *h*_*i*_(*s*_*i*_)’s and *J*_*ij*_(*s*_*i*_, *s*_*j*_) were the fitting parameters of our model. They conform single-site fields and pairwise-couplings that, together, make up the resulting energy score of a given sequence.

Overall, there were 36 single-site (4 bases for each of the 9 sites) and 576 two-site interaction (16 base combinations for 36 site-pairs) parameters that should be estimated in order to fulfill:

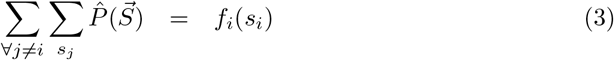

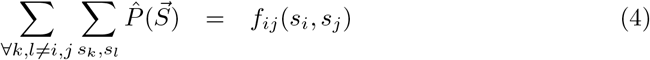

For the sake of convenience, from now on, we will omit the explicit dependency on *s*_*i*_ variables (e.g. *J*_*ij*_(*s*_*i*_, *s*_*j*_) ≡ *J*_*ij*_)

### Fitting procedure

Following [26] we implemented a regularized gradient descent scheme to fit the 612 parameters of our model. In each iteration we generated, using a Metropolis Monte Carlo procedure, an ensemble of 100000 sequences compatible with Eq 1 and the current model parameters. The fitting parameters were then updated according to the following rules:

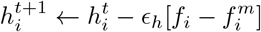

if 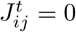

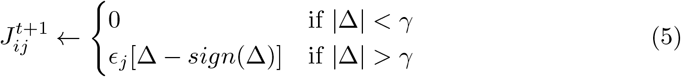

if 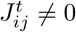

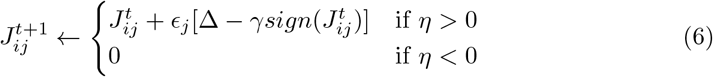

with 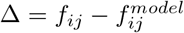 and 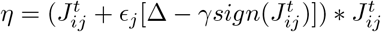.

It can be appreciated from these updating rules that the regularization parameter *γ* prevents non strong-enough coupling constants departing from zero.

### The model

For each analyzed organism, we estimated a family of fitting models. For sequentially decreasing values of *γ* we got models of increasing complexity in a controlled way. With this fitting strategy we could disentangle the hierarchy of minimal sets of coupling constants *J*_*ij*_ necessary to adjust the observed two-site frequencies *f*_*ij*_ at a given accuracy level. We illustrated this point, for the *homo sapiens* case, in Fig 1.

**Fig 1.**
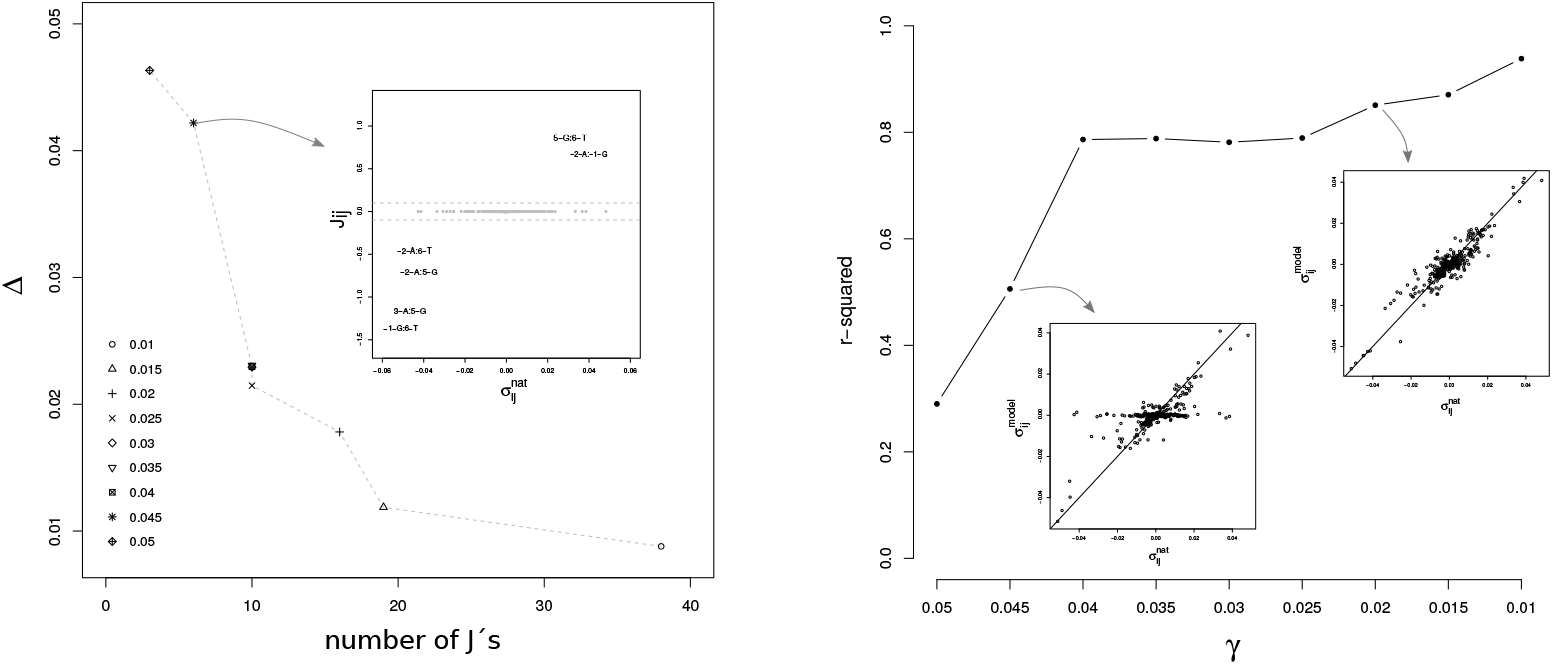
Role of regularization. Left panel: maximum absolute deviation between observed and estimated two site probabilities, Δ = *max*[*abs*(*f*_*ij*_ − *P*_*ij*_)], as a function of the number of the non-zero coupling constants identified at different regularization levels (different symbols, *γ* ∈ [0.01, 0.05]) for human donor sequence models. Inset: coupling constants *J*_*ij*_(*s*_*i*_, *s*_*j*_) as a function of observed two-site correlations 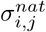 for the *γ* = 0.045 model. Right panel: *r*^2^ correlation coefficient between modeled and observed two-site correlations as a function of the regularization parameter *γ*. Insets: relationship between modeled, 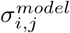 and observed, 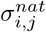, two-site correlations at *γ* = 0.045 and *γ* = 0.02.

In the left panel of Fig 1 we show Δ = *max*[*abs*(*f*_*ij*_ − *P*_*ij*_)] (the maximum absolute deviance between observed, *f*_*ij*_, and estimated, *P*_*ij*_, two-site frequencies) as a function of the number of non-zero coupling parameters. The figure is parametrized by the corresponding regularization *γ* value. It can be observed that a weaker regularization resulted in a more complex model that better fitted the target frequencies. We show in the inset the coupling parameter values, *J*_*ij*_, as a function of the corresponding correlation observed in natural sequences, *σ*_*ij*_ = *f*_*ij*_ − *f*_*i*_*f*_*j*_, for the *γ* = 0.045 case. It can be appreciated that the six non-trivial couplings recognized by this model were associated to the largest two-site correlations present in the data.

At every regularization level the fitting models were able to generate an ensemble of sequences presenting one-site, *P*_*i*_, and two-site frequencies *P*_*ij*_ that accurately reproduced the observed ones (R-squared > 0.99). In the right panel of Fig 1 we show the coefficient of determination (i.e. R-squared) of the regression of model-estimated against observed correlation values. It can be appreciated that with the incremental inclusion of more couplings (at lower regularization levels), the models could reproduce two-site correlations present in natural sequences increasingly well. We found a similar behavior for every analyzed genome (see Sup Fig 1). In particular, it can be seen from this figure that models obtained at *γ* = 0.025 regularization level presented around ten coupling parameters different from zero and achieved a noticeable reduction in Δ. On the other hand, a leveling-off in Δ was consistently observed at *γ* = 0.015.

To better understand the roles and patterns of the fitting parameters, *{h*_*i*_*}* and {*J*_*ij*_}, we show in Fig 2 a graphical summary of the *γ* = 0.025 model for human donor sequences.

**Fig 2.**
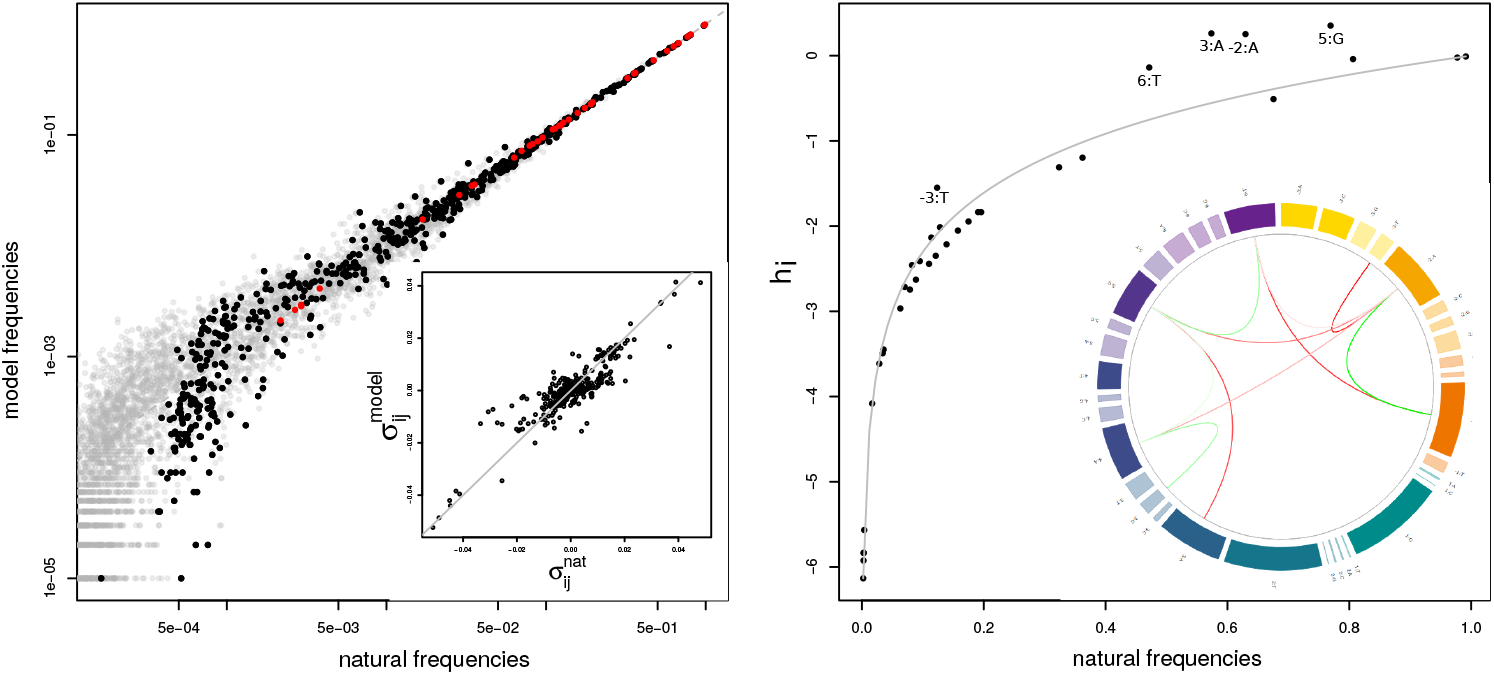
*γ* = 0.025 model frequencies and fitted parameters. Left panel: one-site, two-site and three-site model estimated frequencies (*p*_*i*_, *p*_*ij*_, *p*_*ijk*_) as a function of the corresponding observed frequencies (*f*_*i*_, *f*_*ij*_, *f*_*ijk*_), are depicted as red, black and gray dots respectively. Inset: relationship between model-estimated, 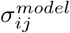, and observed, 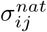 two-site correlations. Right panel: *h*_*i*_(*s*_*i*_) fitting values as a function of the observed one-site frequency, *f*_*i*_(*s*_*i*_). The continuous gray line depicts the independent site model expected relationship 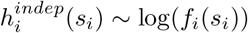. Inset: circos representation of coupling interactions. The 36 circular ring boxes represent the 4 nucleotides per 9-site donor sequence. Warm colors were used for the three exonic sites whereas cold colors for intronic ones. The area of each box is proportional to the nucleotide-site obserevd probability *f*_*i*_(*s*_*i*_). Positive and negative couplings are depicted with green and red lines respectively.

In the left panel of Fig 2 we show the relationship between estimated and observed one-site (red dots) and two-site (black dots) frequencies. The excellent agreement between natural and modeled quantities can be appreciated (R-squared > 0.99 for both cases). The inset of the panel shows the relationship between modeled and observed correlations. At a given level of regularization, correlations were a little bit harder to reproduce (R-squared ∼ 0.84). Noticeably, despite the fitting procedure just considered up to pair-wise observed marginals (i.e. *f*_*i*_ and *f*_*ij*_) we found that the inclusion of a finite-set of pairwise couplings *J*_*ij*_ also allowed our model to reproduce higher-order statistics. In particular three-site probabilities (*f*_*ijk*_, gray dots in Fig2) were very well captured in the simulated ensemble of sequences (R-squared = 0.95).

In the right panel of Fig 2 we show the *h*_*i*_ parameter values as a function of observed frequencies *f*_*i*_. The estimated site-independent approximation for this quantities, *h*_*i*_ ∼ *log*(*f*_*i*_), is depicted as a gray continuous reference line. The inset of the panel is a *circos* graphical representation of the suggested coupling connectivity pattern. The 36 circular ring boxes represent the 4 nucleotides per 9-site donor sequence. Warm colors were used for the three exonic sites, and cool colors for the six intronic ones. Four blocks of different areas, representing the frequency of a given base {*A, C, G, T*}, were included for each site. Positive and negative couplings were depicted with green and red lines respectively.

From the figure it can be observed that many single-site parameters largely deviate from the independent-site approximation. This can be easily understood taking into account the presence of non-zero couplings that were required to reproduce two-site statistics. Positive deviations occur whenever the site is involved in negative interactions and, vice-versa, negative deviations are expected when the site participates in stabilizing interactions. For instance, the single-site fields *h*_5_(*G*) and *h*_6_(*T*), that stabilized the consensus *G* and *T* bases at the last two intronic sites, presented larger values than the independent-site model. This happened because both sites participated in negative interactions with other consensus sites.

### Robustness of inferred coupling patterns

In order to test the robustness of the inferred coupling patterns we focused on the *homo sapiens* case and wondered whether qualitative changes were observed when different subsets of donor sequences were considered. Specifically, we independently obtained *γ* = 0.25 models for: the complete set of annotated donor sequences (502497), a subset of this dataset containing just GT sequences (488939), and two additional subsets of 5’ss expressed in human tissues.

To obtain a maximum entropy model for human 5’ss supported by transcriptomic data we considered the RJunBase resource http://www.rjunbase.org/. This database integrates information about RNA splice linear junctions innormal and cancerous human tissues present in 10,283 RNA-seq samples from The Cancer Genome Atlas (TCGA) and Genotype-tissue Expression (GTEx) portal [27]. We kept only annotated linear junctions of protein-coding genes that were expressed in normal tissues (233961 5’ss) to generate the probabilistic model. A second model was built from a more restrictive set of 114745 joints presenting a median expression level in normal tissue greater than 5.

The four panels included in Fig 3 show the circos diagrams depicting the coupling patterns for the four analyzed models. It can be appreciated that a robust pattern emerge across the four examined datasets.

**Fig 3.**
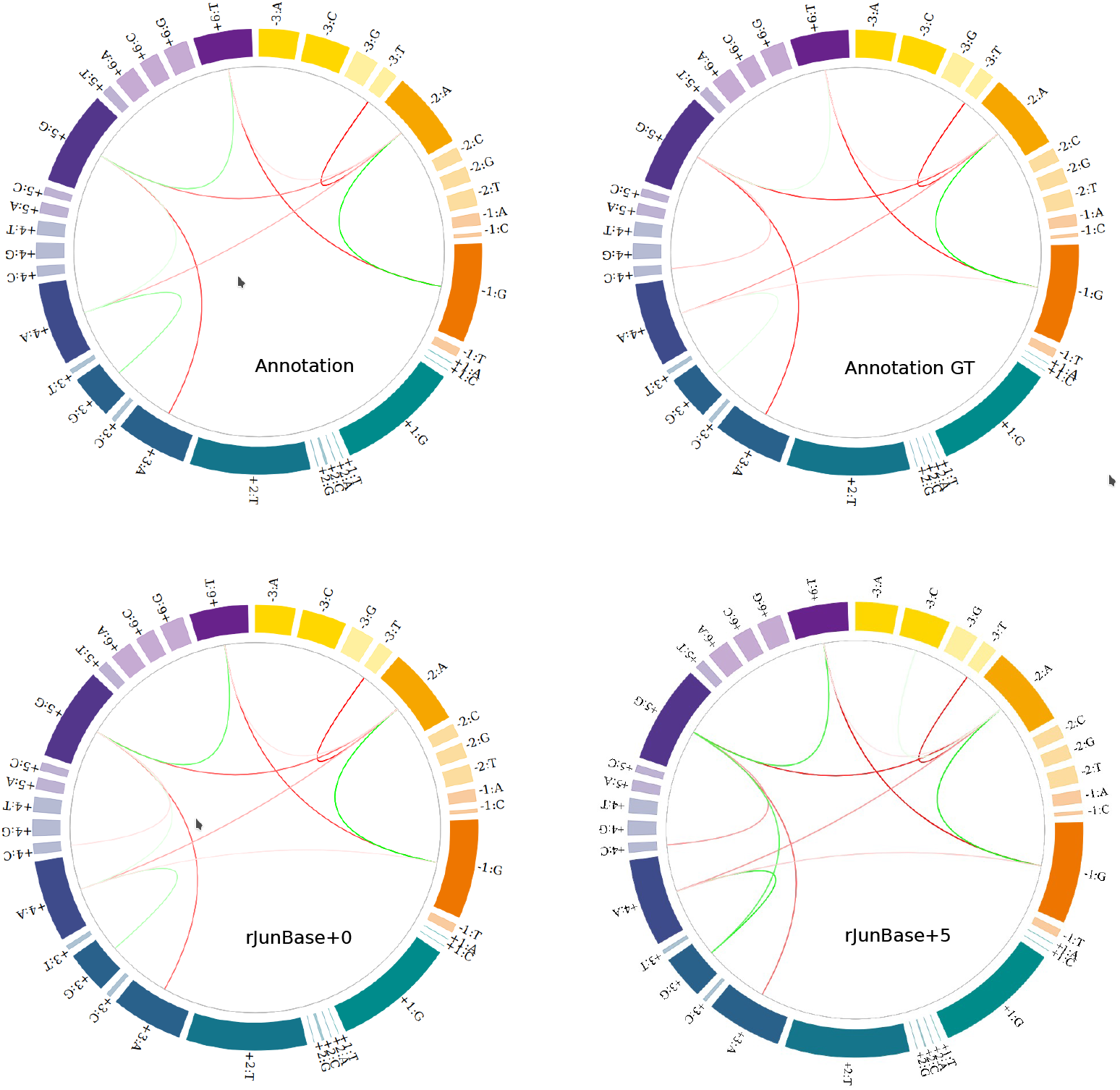
Circos diagrams for coupling patterns estimated for *Homo Sapiens* donor sequences. Patterns for the complete set (502197) and GT-restricted (488939) 5’ss extracted from annotation are shown in the top-left and top-right panels respectively. Patterns corresponding to the complete set of junctions transcriptionally expressed in normal samples according to RJunBase (233961) are shown in the bottom-left panel. The bottom-right panel corresponds to the model inferred from 114745 highly expressed junctions (median count in normal samples larger than 5).

### Time-tree

We got a time-tree for our list of species from timetree.org site (http://www.timetree.org/search/gotimetree, last access December 20, 2021). Two of the thirty analyzed species were not found in the TimeTree database and were replaced by evolutionary close species for the sake of phylogenetic calculations. Specifically, Magnaporthe oryzae (mor) was replaced by Pseudohalonectria lignicola (both from Magnaporthacease family) and Coprinopsis cinerea (cci) was replaced by Coprinopsis lagopus (both from Coprinopsis genus).

### Phyogenetic signals

We considered the Maddison-Slatkin randomization procedure [28] to assess for statistically supported associations between 41 non-zero model coupling parameters and phylogenetic signals. This is a non-parametric bootstrapping approach to generate a distribution of expected values of a test statistic. In our case we considered a parsimony score defined as the number of changes (in parsimony steps) of the binary trait of interest, i.e. the presence/absence state of the analyzed coupling parameter. Specifically, for each coupling parameter a presence/absence binary state was assigned to taxa 10000 times and parsimony scores (Sankoff methodology [29]) were estimated for each random assignment using the functionality implemented in the function *parsimony* of the R-package *phagorn* [30].Bonferroni corrected p-values were estimated from the fraction of random events with parsimony scores equal or larger than the observed value.

### Dendrograms inferred from two-site probabilities

We considered euclidean distances between vectorized upper-triangular matrix of two-site probabilities *P*_*ij*_ of the analyzed organisms. A dendrogram was then produced using the complete agglomeration method.

### Dendrogram comparisons

We made use of the functionality implemented in the R-package *dendextend* [31] to carry out dendrogram comparisons. We found *tanglegrams*, a specific kind of diagrams to visually compare two dendrograms, a very informative tool to qualitatively compare pairs of hierarchical ordinations. In addition we used the function *Bk-permutations* to carry out a bootstrap analysis (10000 permutations) of Fowlkes-Mallows indices to compare partitions induced at different cut-levels from the dendrograms of interest [32]. Given two different partitions of *n* objects, and being *m*_*i,j*_ the number of common elements between the i-th and j-th clusters of the first and second partitions respectively, the Fowlkes-Mallows index is defined as:

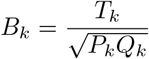

where

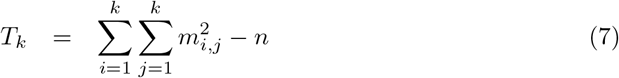

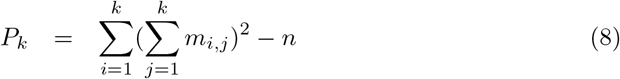

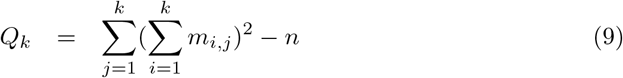

0 ≤*B*_*k*_ ≤1. A higher value for the Fowlkes–Mallows index indicates a greater similarity between the clusters of the two compared partition.

## Results

### Data-driven energetics

According to our estimator of sequence probabilities introduced in Eq 1, the energy function defined in Eq 2 provided a quantitative measure of how often a given sequence can be found along the entire genome. While low energy sequences correspond to the most prevalent splicing sites, high energy ones should be associated to rare and infrequent donor sequences.

Fig 4 shows the frequency distribution of donor sequence energies for the 502197 exon-intron annotated boundaries of the human genome. Ninety percent of sequences presented data-driven energy values laying in the *E*_*d*_ ∈ [4.1, 9.7] energy range. On the low-energy side, the sequence of minimal energy, 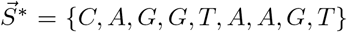, exhibited perfect complementarity to the U1 RNA stretch, presenting an energy value of 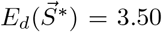. This particular state was found to be the global minimum in the data-driven energy landscape of the entire ensemble of natural sequences (see Sup Fig 3). An ensemble of 2000 randomly generated sequencies was used to get a null reference for our data-driven energy scale. The gray dot and the horizontal black segment in Fig. 4 depict the mean and standard deviations of this null-model distribution (*E*_*d*_ = 21.1 ± 4.6), a proxy for the completely disordered limit.

**Fig 4.**
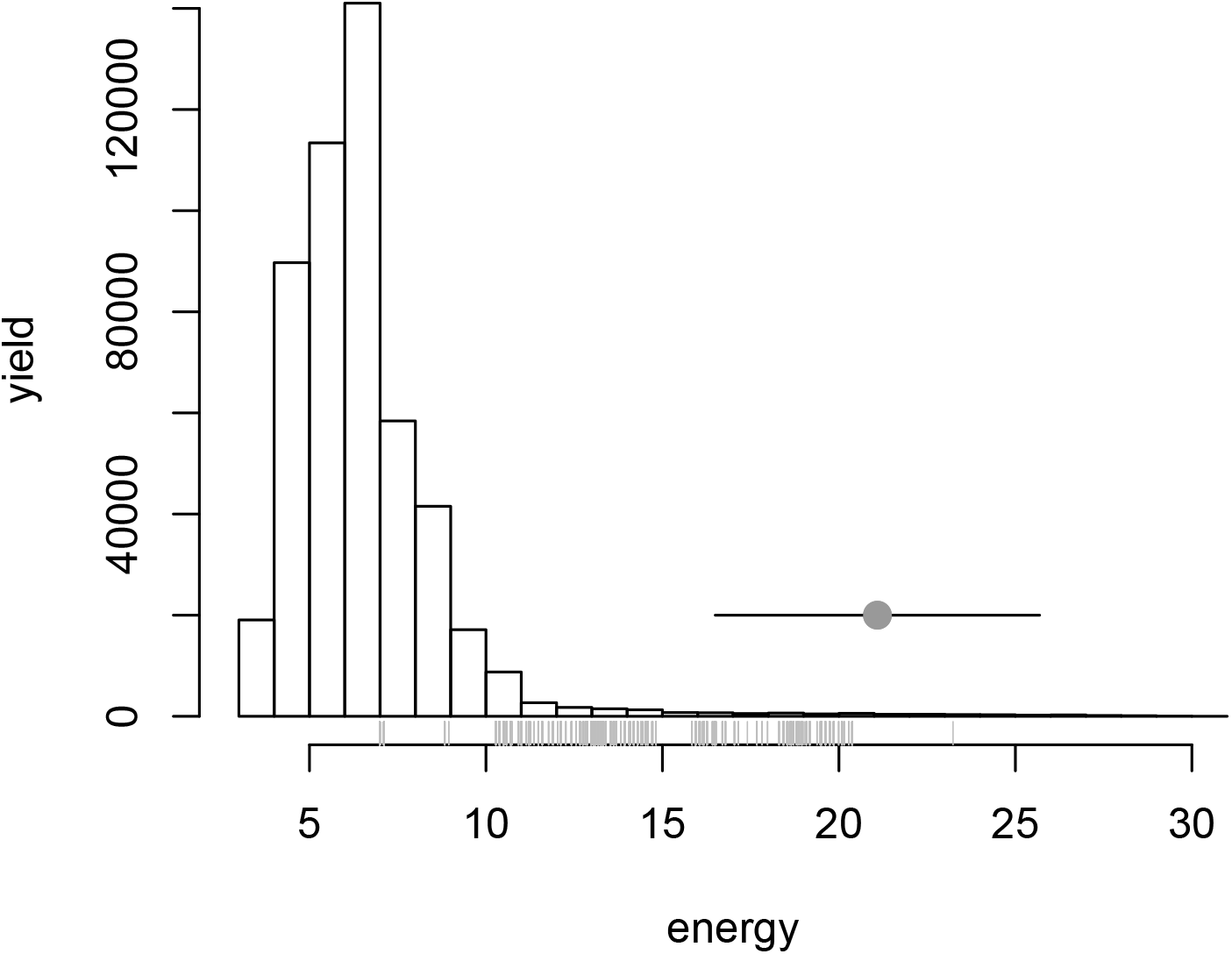
Energetics of human donor sequences (*γ* = 0.025). Distribution of data-driven energy values for the 502197 5’ss sequences of the human genome. Gray ticks were used to mark U12 sequence energies. The gray solid circle corresponds to the mean energy value observed for 2000 random 9-bases length sequences, whereas the black line shows the ±*σ* energy interval of this null distribution.

It can be appreciated that the energy distribution was slightly skewed toward high energy values. In addition, we noticed that high energy sequencies were enriched (gray tick marks in Fig. 4) in the 136 splicing sites that were reported to be targets of the minor U12 spliceosome according to the Intron Annotation and Orthology Database [33]. These findings suggested that correlations in our data-set were mainly driven by statistical regularities of the majoritarian U2 sequences. From our model’s point of view, these U12 donor sites involved rather unusual sequences suggesting that they came from a different statistical distribution. We verified that the main findings (e.g. coupling patterns) and conclusions reported in this manuscript remained unaltered whether these sequences were removed from the training data-set or not and, as we did not have information of the U12 sequences for the other analyzed organisms, we decided to keep them in our data-set.

Fig. 4 also suggested that the data-driven energy (Eq.2) provided a ‘natural’ scale to characterize donor sites, going from completely ordered (perfect match against U1) to completely disordered sequences. Noticeably, the vast majority of exon-intron boundary sequences lied in between these extreme behaviors, in a region of natural variability where recognition is possible but a full binding with the recognition machinery is avoided. In addition, we found that our data-driven characterization correlated with more physically sounded energy scales. Not only the perfect complementary sequence to U1 (i.e. the one that minimized the binding energy) presented the minimal energy value but, more generally, an increasing monotone relationship could be appreciated between our model sequence energies and estimations of the biochemical dimerization energy of 5’ss sequences against the U1 RNA stretch (see Sup Fig 5).

### Conserved coupling patterns

Our fitting procedure allowed us to identify sufficient sets of pairwise coupling interactions that could produce ensembles of sequences compatibles with two-site statistics presented in natural sequences. Table 1 summarizes the average coupling strengths (*γ* = 0.025 models) between consensus (C) and non-consensus (NC) bases found at intronic (I) and exonic (E) sites for each analyzed organism. Despite some lineage specific peculiarities, this comparative analysis allowed us to identify a strong general trend in the pattern of pairwise interactions.

**Table 1.** Conserved patterns (*γ* = 0.025). Mean interactions between different type of sites are shown for different organisms. EC, ENC, IC, and INC stand for exonic-consensus, exonic-non-consensus, intronic-consensus and intronic-non-consensus respectively.

| Species | IC-EC | IC-ENC | INC-EC | INC-ENC | IC-IC | IC-INC | INC-INC | EC-EC | EC-ENC | ENC-ENC |
| --- | --- | --- | --- | --- | --- | --- | --- | --- | --- | --- |
| cne | -0.41 | 0 | 0 | 0 | -0.14 | -0.01 | 0 | 0.88 | 0.13 | -0.13 |
| ani | -0.5 | 0 | 0.03 | 0 | 0.21 | 0.02 | 0 | 0.82 | 0 | -0.15 |
| ncr | -0.68 | 0 | 0 | 0 | 0.26 | 0.03 | 0 | 0.68 | 0 | -0.07 |
| mor | -0.45 | 0 | 0 | 0 | 0.1 | 0 | 0 | 0.87 | 0 | -0.15 |
| cci | -0.59 | 0 | 0.04 | 0 | 0.15 | 0.09 | 0 | 0.7 | 0 | -0.35 |
| ath | -0.39 | 0 | 0.05 | 0 | 0.37 | 0 | 0 | 0.83 | 0 | -0.11 |
| hvu | 0 | 0.09 | 0 | -0.06 | -0.13 | 0.92 | 0.09 | 0 | -0.16 | -0.29 |
| mtr | -0.3 | 0 | 0 | 0 | 0.1 | 0 | 0 | 0.95 | 0 | 0 |
| osa | -0.24 | 0 | 0.01 | 0 | 0.23 | 0.05 | -0.04 | 0.94 | 0 | 0 |
| ppa | -0.77 | 0 | -0.01 | 0 | -0.02 | -0.01 | -0.03 | 0.64 | 0 | -0.08 |
| ptri | -0.42 | 0 | 0 | 0 | -0.02 | 0 | 0 | 0.91 | 0 | 0 |
| sly | -0.36 | 0 | -0.01 | 0 | 0.21 | 0 | 0 | 0.91 | 0 | -0.07 |
| vvi | -0.37 | 0 | 0 | 0 | -0.02 | 0 | 0 | 0.93 | 0 | -0.04 |
| apl | -0.4 | 0 | 0 | 0 | 0.29 | 0.01 | 0 | 0.87 | 0 | 0 |
| bta | -0.28 | 0 | 0 | 0 | 0.57 | 0 | 0 | 0.78 | 0 | 0 |
| clu | -0.3 | 0 | 0 | 0 | 0.52 | 0.03 | 0 | 0.79 | 0 | 0 |
| dre | -0.45 | 0 | 0 | 0 | 0.23 | 0 | -0.02 | 0.86 | 0 | 0 |
| eca | -0.37 | 0 | 0 | 0 | 0.48 | 0 | 0 | 0.8 | 0 | 0 |
| ggo | -0.18 | 0 | 0 | 0 | 0.59 | -0.01 | 0 | 0.78 | -0.1 | 0 |
| hsa | -0.48 | 0 | 0 | 0 | 0.14 | -0.01 | 0 | 0.86 | 0 | 0 |
| mdo | -0.22 | 0 | 0 | 0 | 0.67 | 0 | 0 | 0.69 | -0.15 | 0 |
| mmu | -0.51 | 0 | 0 | 0 | 0.08 | 0 | 0 | 0.86 | -0.06 | 0 |
| oan | -0.19 | 0 | 0 | 0 | 0.81 | -0.06 | 0.03 | 0.55 | 0 | 0 |
| ocu | -0.24 | 0 | 0 | 0 | 0.61 | 0 | 0 | 0.76 | 0 | 0 |
| sha | -0.33 | 0 | 0 | 0 | 0.6 | 0.01 | 0 | 0.73 | -0.05 | 0 |
| ssa | 0.14 | 0 | -0.06 | 0 | 0.74 | 0.01 | 0.02 | 0.63 | -0.16 | 0 |
| ssc | -0.14 | 0 | 0 | 0 | 0.7 | -0.04 | 0 | 0.7 | -0.07 | 0 |
| xtr | 0.11 | 0 | -0.04 | 0 | 0.56 | -0.02 | 0.03 | 0.81 | -0.13 | -0.04 |
| dme | -0.89 | 0 | 0 | 0 | -0.23 | -0.04 | 0 | 0.38 | 0 | -0.08 |
| cel | -0.9 | 0 | 0.05 | 0 | -0.21 | -0.01 | 0 | -0.35 | 0 | -0.13 |

Stabilizing positive interactions could be recognized for consensus bases inside exonic (EC-EC) and intronic portions (IC-IC) of donors sites. Noticeably, a clear negative coupling between intronic and exonic consensus sites (IC-EC column in Table 1) could also be recognized across all the analyzed species. This last finding, suggesting that simultaneous co-apparition of consensus sites both, at the intronic and exonic parts of donor sequences, was statistically less preferred, extended already presented evidence of negative epistatic signals found in mammals donor sequences [18, 19]. These results were also observed for more complex *γ* = 0.015 models (see Supplementary Table 2).

### Divergent coupling signals

Despite the general high degree of conservation of the spliceosome, single site frequencies were shown to display a non-trivial degree of specificity in eukaryotes [15, 34]. Noticeably we found that two-site marginal probabilities captured by our model also exhibited this behavior. In fact, the hierarchical structure obtained from *P*_*ij*_ probabilities was identical to the one inferred from logo motifs, i.e. one site statistics *P*_*i*_ (see Sup Fig 6).

In the left panel of Fig.5 we show a tanglegram. The left-most dendrogram was built considering two-site probabilities *P*_*ij*_ for *γ* = 0.025 models whereas the second one was the time-tree inferred from phylogenetic signals.

**Fig 5.**
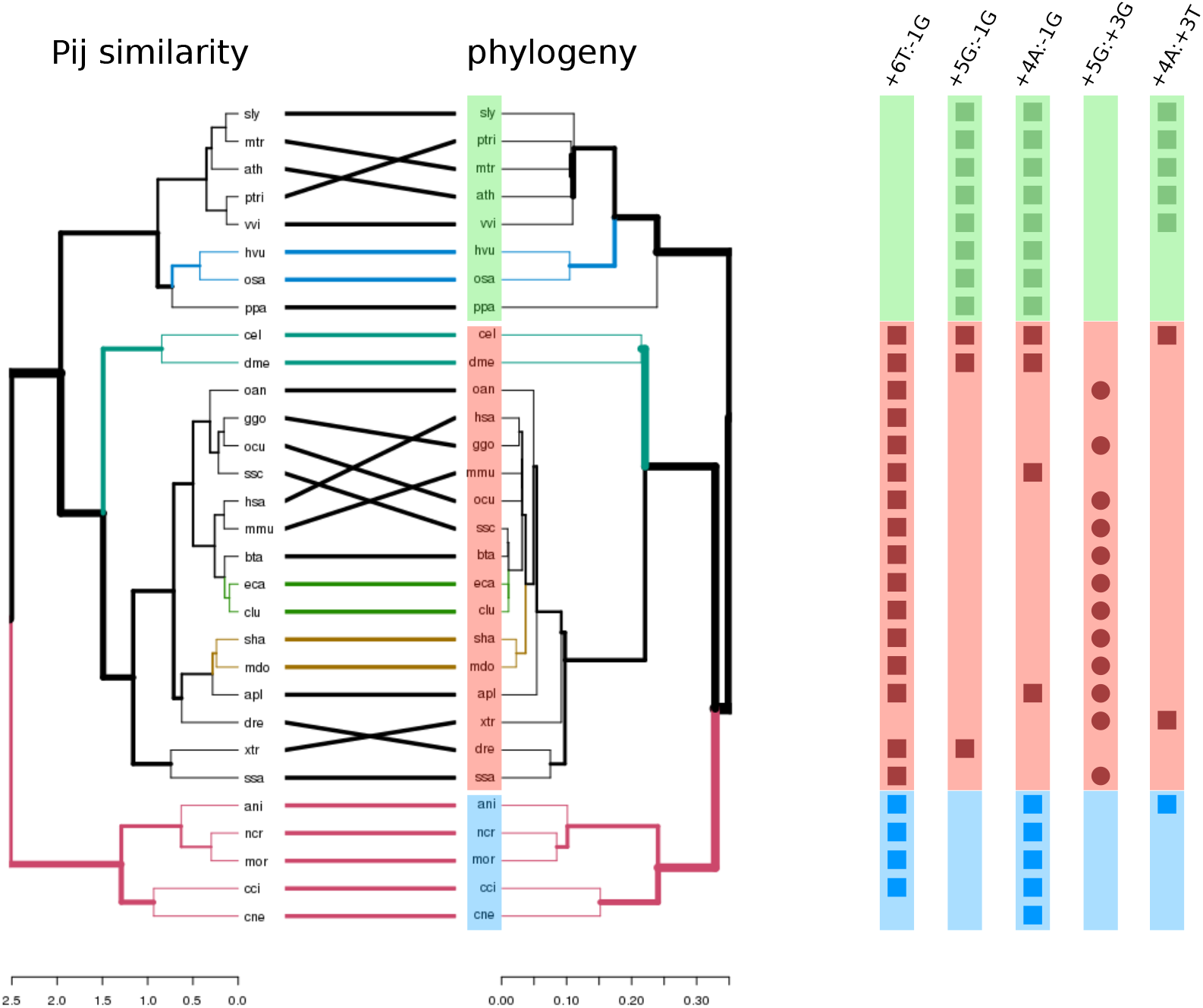
Divergent coupling patterns. **A**, Tanglegram layout for the *P*_*ij*_-induced dendrogram (left) and the pyhilogenetic tree (right) of the analized organisms. Green, red and blue are used to highlight plant, animals and fungi respectively. **B**, Presence/absence matrix for the seven statistically significant model interaction parameters (see text). Square and circle shapes were used for negative and positive interactions respectively.

It can be appreciated from the figure that the ordering structure from two-site statistics *P*_*ij*_ displayed strong concordance with the underlying phylogenetic relationships between the involved organisms. In particular a clear separation between plants, animal and fungi could be observed. Several quantitative figure of merits supported this finding. For instance, a cophenetic correlation value of 0.9 was found between both dendrograms, and Fowlkes-Mallows indexes were found to be statistically significant (*p*_*v*_ < 10^−4^) for almost the entire range of k-cluster groups (2 < *k* < 29) (see Sup Fig 4). Additionally, an entanglement value of 0.01 suggested a high quality tanglegram layout (see Material and Methods).

### Coupling parameters and phylogenetic signals

Panel A in Figure 5 suggested that two-site statistics captured by our model contained information compatible with phylogenetic signals. We wondered if this signal was also caught at the level of the coupling parameters highlighted by our maximum entropy model.

For each one of the 41 non-zero *J*_*ij*_ identified in any of the analyzed organisms, we performed a Maddison-Slatkin procedure as a test for phylogenetic signal (see Material and Methods). We found five statistically significant associations and reported the corresponding results in Table 2. The number of observed evolutionary steps for the analyzed character, inferred using the Sankoff parsimony methodology [35], are shown in column SM.obs. The minimum, median and maximum number of evolutionary steps detected in 10000 bootstrapped samples, and Bonferroni corrected p-values were reported in columns SM.null and SM.pv respectively. The corresponding presence/absence patterns of these interactions are displayed in Panel B of Figure 5. For the sake of clarity we used a simplified notation for the coupling parameters. For instance -1G:+6T denoted the *J*_−1,+6_(*G, T*) coupling parameter.

**Table 2.** Phylogenetic signals associated to coupling parameters. Statistically significant model parameters for Maddison-Slatkin tests are reported in rows. For each coupling parameter (first column) we specified in how many plants, animals and fungi the interaction was detected (second, third and fourth columns). The observed number of Sankoff inferred evolutionary steps is reported in the fifth column. Minimum, median and maximum values for this quantity for bootstrapped samples are reported in the sixth column (comma separated values). Bonferroni corrected pvalues were included in the last column of the table.

| Parameter | plants | animals | fungi | MS.obs | MS.null | MS.pvD |
| --- | --- | --- | --- | --- | --- | --- |
| -1G:+6T | 0 | 16 | 4 | 3 | 3,9,10 | 4.1e-3 |
| -1G:+5G | 8 | 3 | 0 | 3 | 4,9,11 | 0 |
| -1G:+4A | 8 | 4 | 5 | 3 | 3,10,13 | 4.1e-3 |
| +3G:+5G | 0 | 12 | 0 | 4 | 4,10,12 | 4.1e-3 |
| +3T:+4A | 5 | 2 | 1 | 3 | 3,7,7 | 5.0e-2 |

As expected, it can be appreciated from this figure that the identified coupling parameters were selectively present in plants, animals or fungi. The first three entries of Table 2 corresponded to negative interactions between intronic and exonic consensus nucleotides that involved the -1G exonic element whith the last three intronic positions. Interaction -1G:+6T was found to be a trait for animals-and-fungi, -1G:+5G was mainly present in plants. The -1G:+4A coupling, on the other hand, could be considered a trait for plants and fungi.

The fourth entry in Table 2 corresponded to a positive intron-intron interaction between consensus nucleotides (+3G:+5G) that appeared in 70% of the analyzed metazoans with no plants or fungi displaying it at all. Finally the +3T:+4A negative coupling was found in 60% of the analyzed plant genomes.

Our findings suggested that non-trivial phylogenetic information was actually present in two-site correlations that, in our model, were linked to statistical coupling patterns. In particular the strongest phylogenetic signals were reported for coupling parameters that involved a negative effective interaction between a consensus nucleotide in the last exonic position and consensus nucleotides located at the last three intronic sites.

### Consolidated models

In order to further investigate the specificity of coupling signatures, we wondered if characteristic coupling patterns could be associated to animals, plants and fungi. We consolidated donor sequences performing a uniform and balanced sampling of 800000 splicing sequences for the 17 metazoans, 800000 splicing sequences for the 8 plants, and of 161547 sequences for the 5 fungi organisms. We then estimated maximum entropy models for each data-set (see Methods) in order to get representative coupling patterns for each analyzed group.

The resulting coupling diagrams are shown in Fig.6. In the first row of this panel we included results for *γ* = 0.025 models for plants, animals and fungi in the first, second and third columns respectively. In this case, the rather strong regularization level served to highlight the main coupling patterns involved in the establishment of the most significant two-site correlations observed in each data-set (see Fig. 2 inset).

**Fig 6.**
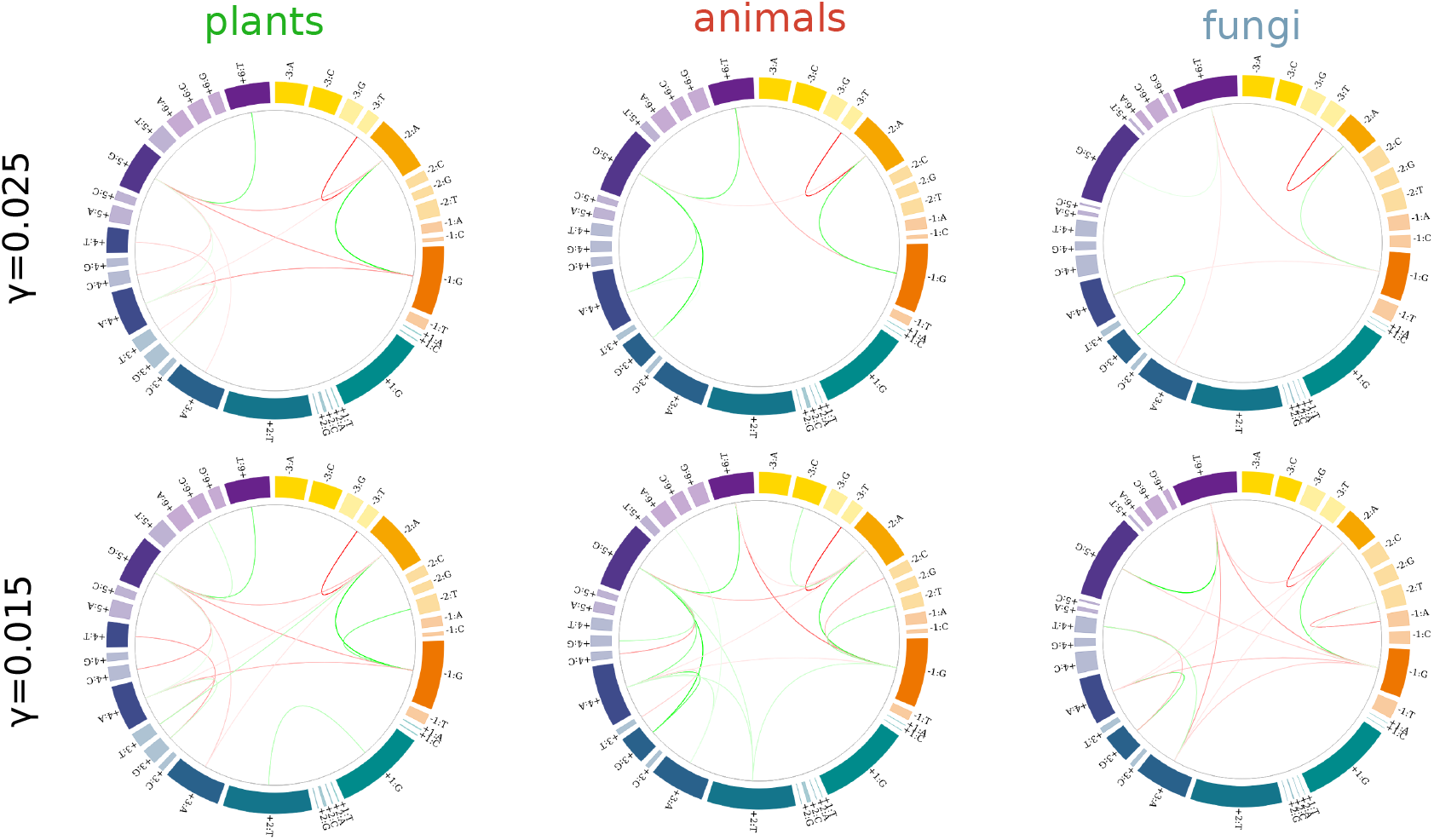
Pairwise interaction patterns. Circos diagrams for coupling parameters identified for plants, animals and fungi donor sequences are shown in columns. *γ* = 0.025 (first row) and *γ* = 0.015 (second row).

Many of the coupling parameters identified in previous sections could be recognized. For instance, positive couplings between consensus sites at the intronic and the exonic part of the splicing junction (e.g. -2A:-1G or +5G:+6G) were present in plants, animals and fungi. Another ubiquitous interaction was the negative coupling between the non-consensus and consensus exonic nucleotides -2A:-3T.

Noticeably we also found patterns consistent with the divergent behavior uncovered by the Maddison-Slatkin analysis. A strong negative coupling was observed between +6T and -1G in the metazoan and fungi groups, that was replaced by a negative interaction between +5G and -1G nucleotides in the plant group (in the next section we further elaborated on this point). On the other hand, the negative coupling -1G:+4A was detected for plants and fungi, but not for metazoans, whereas the positive coupling +3G:+5G was a signature only detected for this last group. All of these patterns persisted in more complex models with relaxing regularization levels, like the one obtained at *γ* = 0.015 shown in the second row of Fig 5-B.

## Discussion

The molecular recognition of 5’ splicing sites faces numerous challenges. First, with the exception of the first GT intronic nucleotides, natural 5’ss sequences are highly degenerate. Secondly, recognition occurs at different stages of the splicing cycle from various complexes that bind totally or partially to the 5’ss sequence. Thirdly, not only snRNAs are involved in recognition, but also multiple proteins present in the complexes are important for stabilizing the binding. Finally, splicing takes place in a genomic context in which factors such as gene structure and the presence of cis and/or trans signals can exert a significant influence on the fine regulation of each particular splicing site. In this work we aimed to mine conserved and divergent signals in splicing donor sequences. The rationale of our approach was that underlying statistical patterns in donor sequences composition should reflect this complex scenario.

Our entropy maximization strategy allowed us to recapitulate in a unified framework previous results and to gain new insights about regularities embedded in donor splicing sequences. For instance, the data-driven energy scale *E*_*d*_ (Eq. 2), naturally accommodated the idea behind the SD-Score (defined as the logarithm of the frequency of a donor sequence) introduced by Sahashi and collaborators to predict the splicing outcomes observed in artificially designed minigenes [18]. *E*_*d*_ also correlated with estimated dimerization energies against U1 snRNA (see Sup Fig 5). Albeit that there are a large number of cis and trans elements that contribute to the regulation of the splicing process, this finding suggested that *E*_*d*_ by itself might reflect at some degree the strength of the donor site. In this sense, *E*_*d*_ not only served, by definition, to quantify how much a given sequence was represented along a genome, but at the same time acted as a meaningful scale to measure the degree of complementarity to the spliceosome machinery. The majority of naturally occurring sequences presented intermediate *E*_*d*_ values (Fig. 4). The fact that a 9-nucleotide length perfect match was avoided agreed with previous observations stating that a minimal 5-6 Watson-Crick pairs were required for splicing site recognition, but too much pairing (> 7 bases) would be detrimental [17]. This loose binding might favor splicing reaction processivity and could favor scenarios where the effective binding could be regulated by third players.

With the aid of our model we could also identify quite general two-site interaction patterns that suggested that this free-energy deficit was rooted in a non-trivial spatial distribution of matching pairs along splicing site sequences. On one hand, despite some degree of specificity, couplings between consensus sites inside exonic and intronic parts of donor sequences were biased toward positive values (columns 5th and 8th of Table 1). On the other, negative interactions were found between consensus nucleotides laying at different sides of the exon-intron boundary (first column of Table 1). These results extended previous observations obtained for human and mouse donor sequences [17–19] and supported the idea that a high complementarity level is alternatively favored either at the exonic or the intronic part of the splicing site.

Two-site couplings reflected that nucleotide identities at different positions of 5’ splicing sites were not independent. Albeit this observation has been reported in the past we showed, as far as we know for the first time, that joint probabilities of nucleotide pairs carried biologically meaningful information in the sense that dendrograms inferred from them closely followed phylogenetic relationships between the analyzed organisms (see Fig 5). Donor splicing sequences constitute recognition sites not only for U1 snRNA in the early spliceosome complex, but also for U6 and U5 in the pre-catalytic reaction step. Subtle compensation mechanisms could then be expected to occur to assure splicing fidelity even in the presence of large site variability. In this conext, our findings suggested that the uncovered statistical regularities might echoed evolutionary divergent processes linked to structural specificities of the splicing machinery used to define exon-intron boundaries [34].

Our results also suggested that phylogenetic signals could be captured by the coupling parameters of our model. These associations proved to be robust in the sense that they were detected for strong and mild regularized models, when coupling patterns were analyzed for each organism separately, or when splicing donor sequences were consolidated into plants, animals and fungi groups.

Many of the interactions detected for the different organisms analyzed in this contribution were already reported in the literature in the context of narrower studies focused on human, mouse or some other mammalian genomes [13, 16–20]. For instance, positive couplings between intronic +4:+5 and +5:+6 position-pairs, and negative interactions between -2:+5 were readily captured by our model. Interestingly we found a -1:+5 negative coupling for plants but, unlike previous studies of human and mammals organisms, we did not detect it for metazoans. This interaction involved the informative -1 and +5 positions (see Sup Fig 9) for which a negative interaction was reported in previous studies like the work of Yeo and Burge, who analyzed 12700 introns of 1821 non-redundant transcripts [20], or the contribution of Carmel and collaborators, who inferred the -1G:+5G compensatory relationship from a comparative analysis of 8869 human-mouse homolog exons [17]. Interestingly, in a recent contribution Wong and collaborators quantifyed the activity of artificially engineered 32,768 unique 5′ ss sequence in three different genomic contexts and found a positive epistatic interaction between these sites [12]. In our case we relayed our comparative analysis in annotated splicing sequences and could not captured the aforementioned interaction neither for the 800000 metazoan splicing sequences nor for the 502197 sequences of human splicing donors. In fact, right from start, we found a fairly low absolute correlation value between -1G and +5G nucleotide frequency of apparition in our metazoan 5’splicing sequence dataset (this behavior was observed regardless the complete set of annotated 5’ splicing sequence was considered, or when we restricted the calculation to donor sequences presenting the canonical GT nucleotides at the intron’s beginning, see Sup Fig 7). According to our model, the compensatory behavior reported between the last exonic position and intronic sites was implemented by an interwoven set of interactions that, for one hand stabilized +5G:+6T and -1G:-2A consensus nucleotides, and for the other negatively coupled -2A:+5G and -1G:+6T sites (see Fig. 6). Noticeably, using data from *homo sapiens*, we showed that these results were obtained not only in models inferred from annotated donor sequences, but also when we considered human 5’ss supported by transcriptional data (see Sup Mat).

These intermingled interaction pattern was consistent with recent observations of Artemyeva and Porter who reported that the base pair composition at positions -1 and -2 were significatively altered based on the absence/presence status of a +5G nucleotide (see Fig 4 in [11]). In addition, the relevance of position +6 was already noticed by Carmel and collaborators in connection with splicing aberrations leading to familial dysautonomia [17]. In that work, the authors provided experimental evidence showing that a base pair of position − 1 rescued an aberrant splicing of the 20th intron of the IKBKAP gene caused by a mispair of position +6 with U1 snRNA.

In our study we found a similar compensatory setup between intronic (+5 and +6) and exonic (−2 and -1) positions in plants, animals and fungi. Specifically we detected for plants a -1G:+5G negative coupling that replaced the -1G:+6T interaction observed in metazoans (see Fig. 6). Noticeably the involved two-sites interactions affected intronic positions that had fairly low information content in plants (0.27 bits and 0.21 bits for positions +5 and +6 respectively, see Sup Fig 9) suggesting that the detected network of pair-wise interactions might play a major role for these organisms to deal with sequence variability.

Several differences were already highlighted between plants and animals in regard to splicing mechanisms. Arguably, the most straightforward ones involve large differences reported in typical intron lengths and the prevalence of different kind of splicing events: exon skipping for animals and intron retention for plants [5,36]. Many recent contributions highlighted not only gene-architectural but also functional differences between alternative splicing taking place in animals and plants [37–40]. For instance, despite the spliceosome in plants have not been isolated yet, the number of splicing factor identified in *Arabidopsis thaliana* nearly double the ones observed in humans [41]. In addition, the prevalence of intron retention events coupled to NMD transcript degradation and nuclear sequestration suggested that, differently from animal organisms, splicing in plants could play a major functional regulatory role, closely related to stress response, apart from expanding proteomic diversity. [5, 36, 42–44]. In this context, our results highlighted significant differences in two-site interactions involving donor site nucleotide positions relevant at functional and evolutionary levels. Based on our findings we believe that the specific connection of these differences with distinctive mechanistic features of splicing processes in plant and animal organisms deserves to be further investigated.

## Conclusion

In this work we followed a maximum entropy program to obtain regularized probabilistic generative models of donor sequences for 30 different eukaryote organisms.

Our approach allowed us to introduce a data-driven energy scale that reflected the abundance of a given sequence along a specific genome. This energy statistic provided a sensible scale to characterize a given sequence in connection with the idea of completely ordered and/or completely disordered sequence states.

In our work we also showed that joint di-nucleotide probabilities of donor sequences carry lineage information. With the aid of our models, we could identify minimal sets of coupling patterns that could produce, at a given regularization level, observed two-site frequencies in donor sequences. The analysis of these interactions across organisms allowed us to identify specific two-site coupling patterns differentiating plants, animals and fungi. This sequence composition signature was embedded in two-site interactions involving the last nucleotides of the intronic part of donor sequences, which suggested that they could be related to taxon-specific features of the early and pre-catalytic spliceosome.

## Supporting information

**Sup Fig 1.**
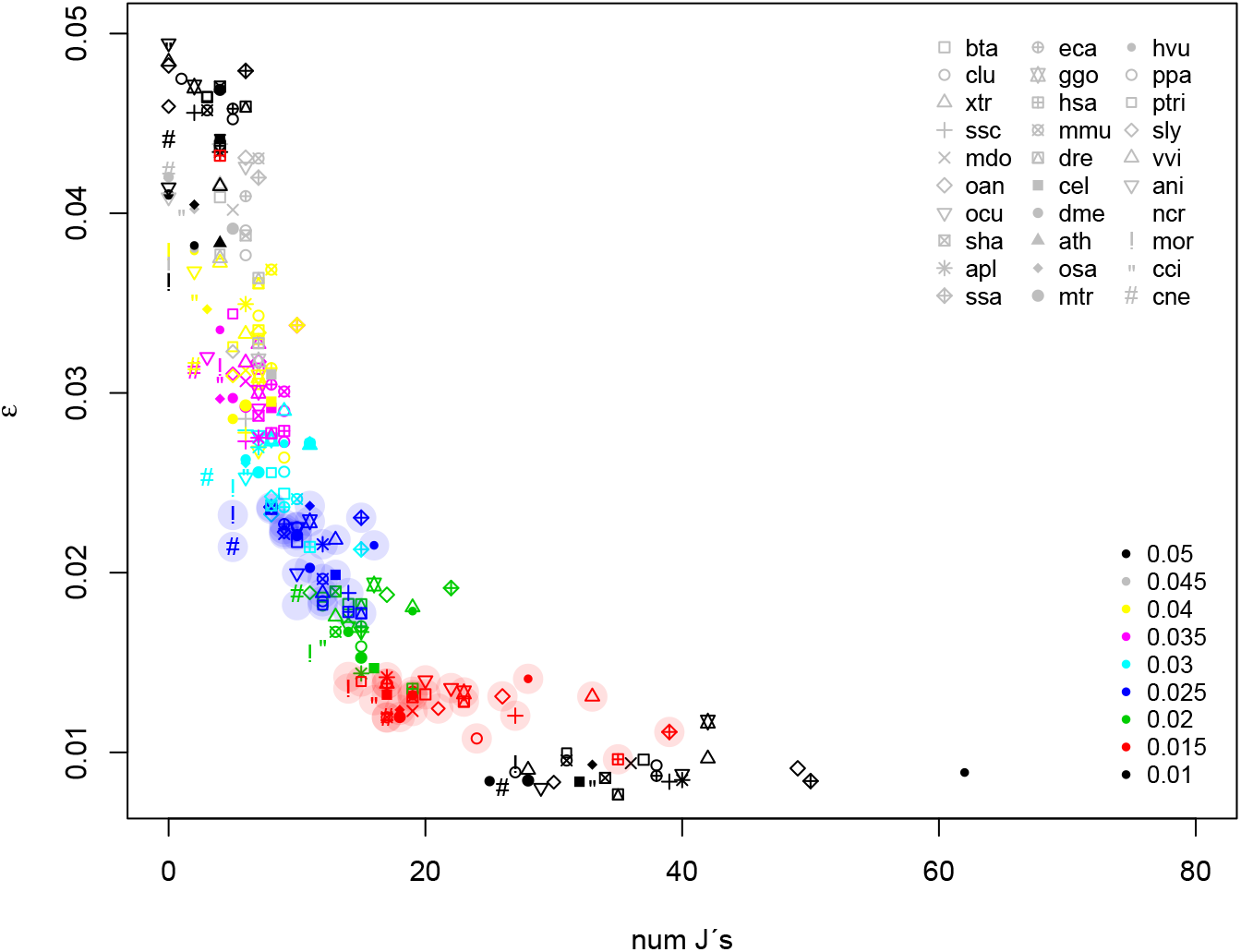
Model convergence. Maximum absolute deviance between observed and estimated two site probabilities, Δ = *max*[*abs*(*f*_*ij*_ − *P*_*ij*_)], as a function of the number of the non-zero coupling constants identified at different regularization levels (different colors, *γ* ∈ [0.01, 0.05]). Results for different analyzed organisms are depicted with different symbols (see legend). Shaded symbols highlight *γ* = 0.025 (blue) and *γ* = 0.015 (red) models respectively.

**Sup Fig 2.**
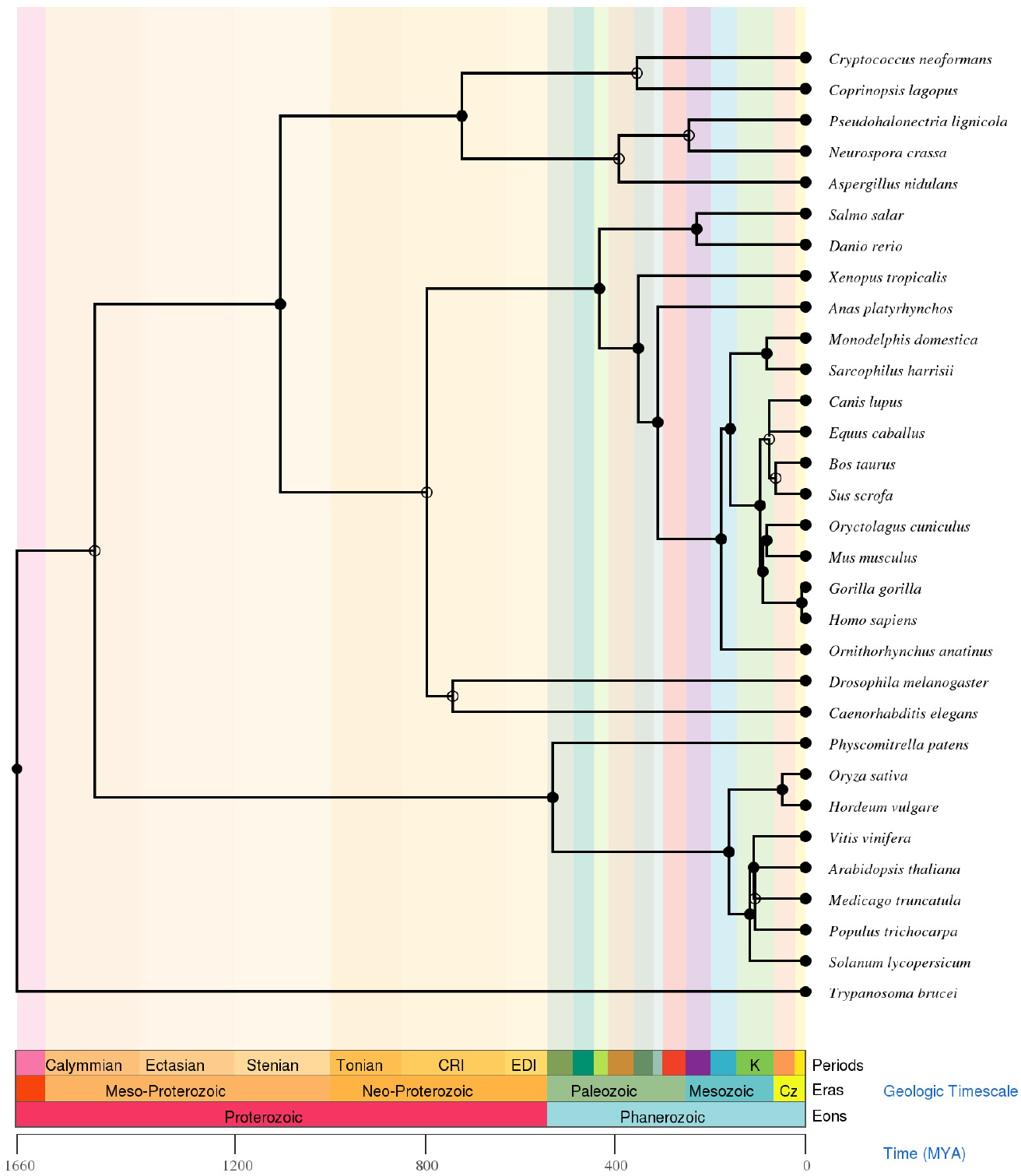
Phylogenetic tree. Time-tree inferred from phylogenetic signals for the 30 analyzed eukaryote organisms. Generated at the timetree.org site (http://www.timetree.org/searchtimetree, December 20, 2021).

**Sup Fig 3.**
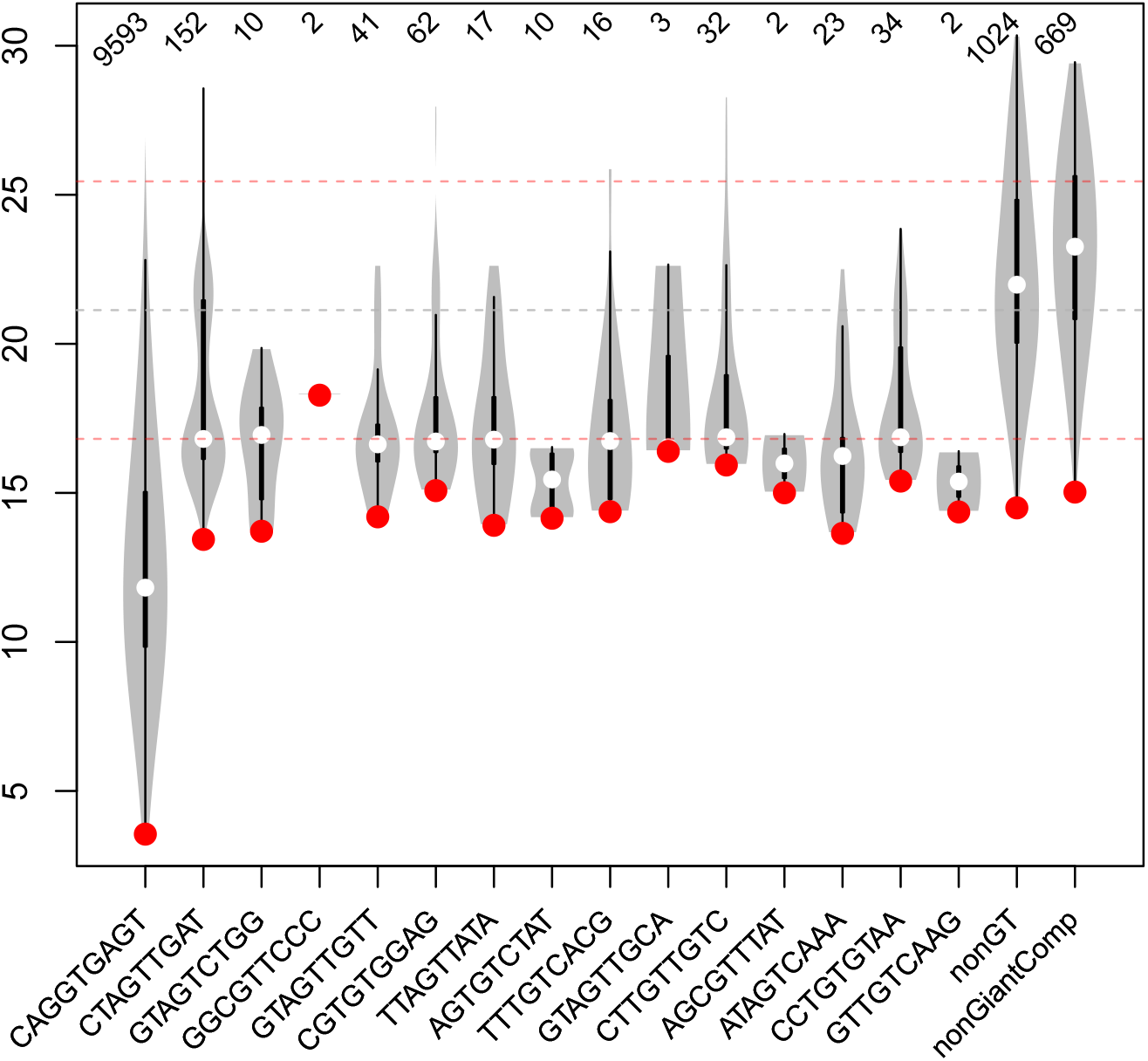
Energy landscape. To further characterize the energy landscape of donor sequences, we analized the *γ* = 0.025 model for human donor sequences considering a network graph approach. Representing individual sequences with nodes, we connected two of them if they were one mutation away (i.e. Hamming distance of 1). The edge started at the node with higher energy and ended in the lower energy one. In this directed network, we found 243 local-minima sequences associated to nodes with no outwards edges (i.e. null out-degree). For each one of these attractor-nodes we could estimate the in-ward connected component, i.e. the set of nodes that could reach a given local minimum-energy node in a finite number of greedy steps. These sets of nodes were a proxy of the extent of the basin of attraction of a given attractor sequence.

Violin plots for the energy distribution of sequences belonging to different basins of attraction are shown in this figure. Each violin plot shows the lowest and highest energies within the corresponding basin and summarizes the overall energy distribution. The first 15 boxplots depict the energy distribution of the basins of the 15 attractor sequences presenting a GT di-nucleotide at the first two intronic positions. The two last boxplots correspond to the energy distribution of non-GT attractor sequences’s basins, and sequences that did not belong to the giant component of the graph respectively. Red dot highlight the attractor (i.e. minimal) energy of each set. The corresponding sequence is included as an x-axis label. Top label show the number of sequences in each basin.

The first boxplot in Fig Sup Fig 3 corresponds to the huge main basis of attraction that contained 9593 sequences. This number represented 80% of the complete set of 5’ exon-intron boundaries of the *H*.*sa* genome, and 96% of the entire set of GT 5’ss.

Noticeably, the associated attractor state sequence 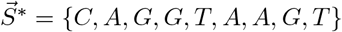 was the perfect matching sequence of U1 smRNA and presented an energy value 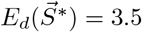. It can be seen from the figure that the basins of attraction of the rest of the local minima presented much higher energy values in much shallower energy wells. Among these secondary local-minima, there were just 14 attractors presenting GT as first intron nucleotides.

These results suggested that in terms of the model data-driven energy our system presented a rather well-defined global minimum state laying inside a wide energy well.

**Sup Fig 4.**
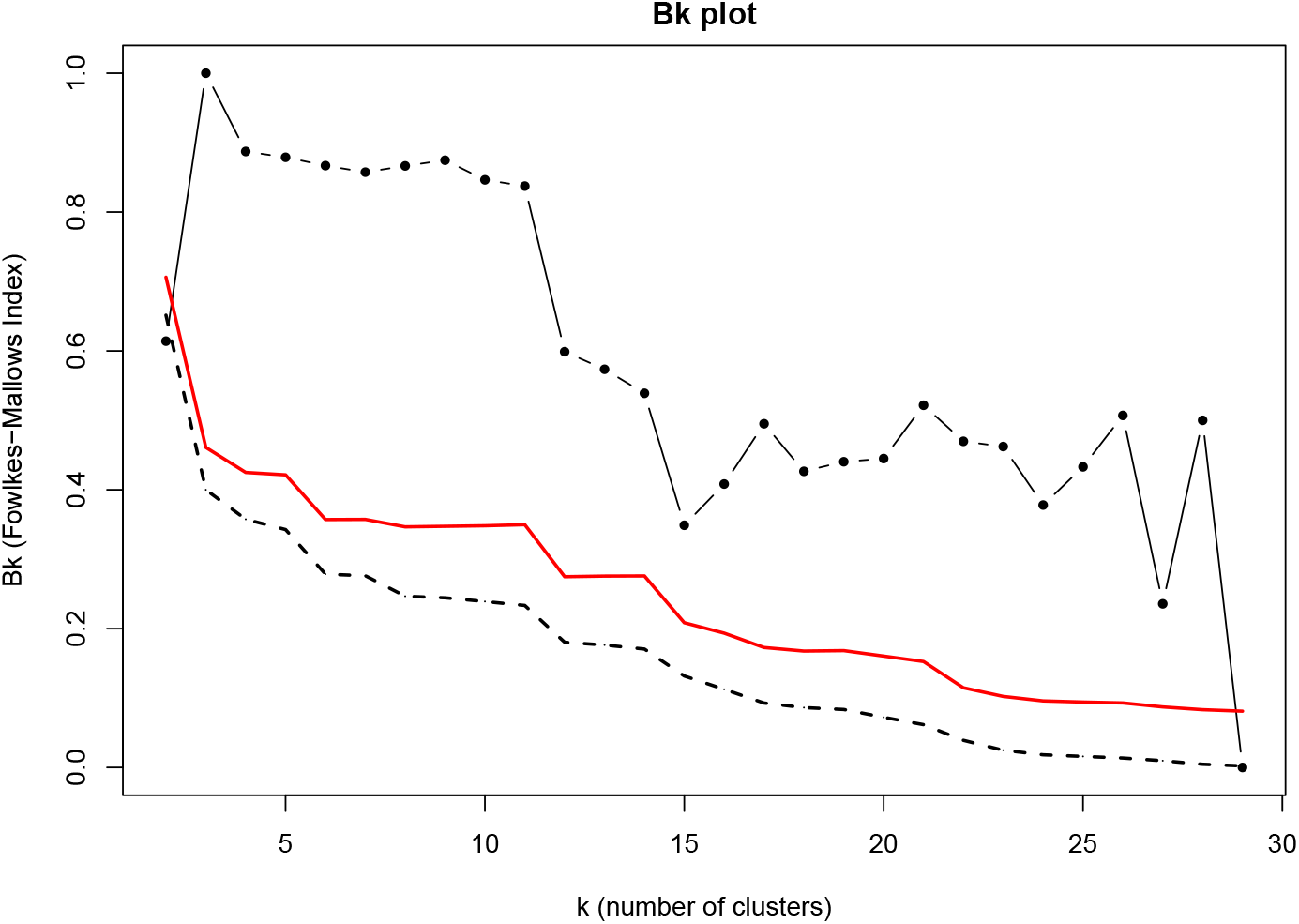
Fowlkes-Mallows analysis. K-cuts partitions were produced for phylogenetics and *P*_*ij*_-induced dendrograms. For each k value the Fowlkes-Mallows clustering similarity index, *B*_*k*_, was estimated along with 10000 null-model values obtained by label shuffling. The dotted black line corresponds to observed *B*_*k*_ values as a function of number of clusters *k*. The expected values under the null hypothesis of no relation between clusterings are shown as a black dashed line. The full red line depicts the 95% one-side rejection region based on the asymptotic distribution of *B*_*k*_ values. All calculations were done using the *Bk_plot* function of the *dendextend* R-package [31].

**Sup Fig 5.**
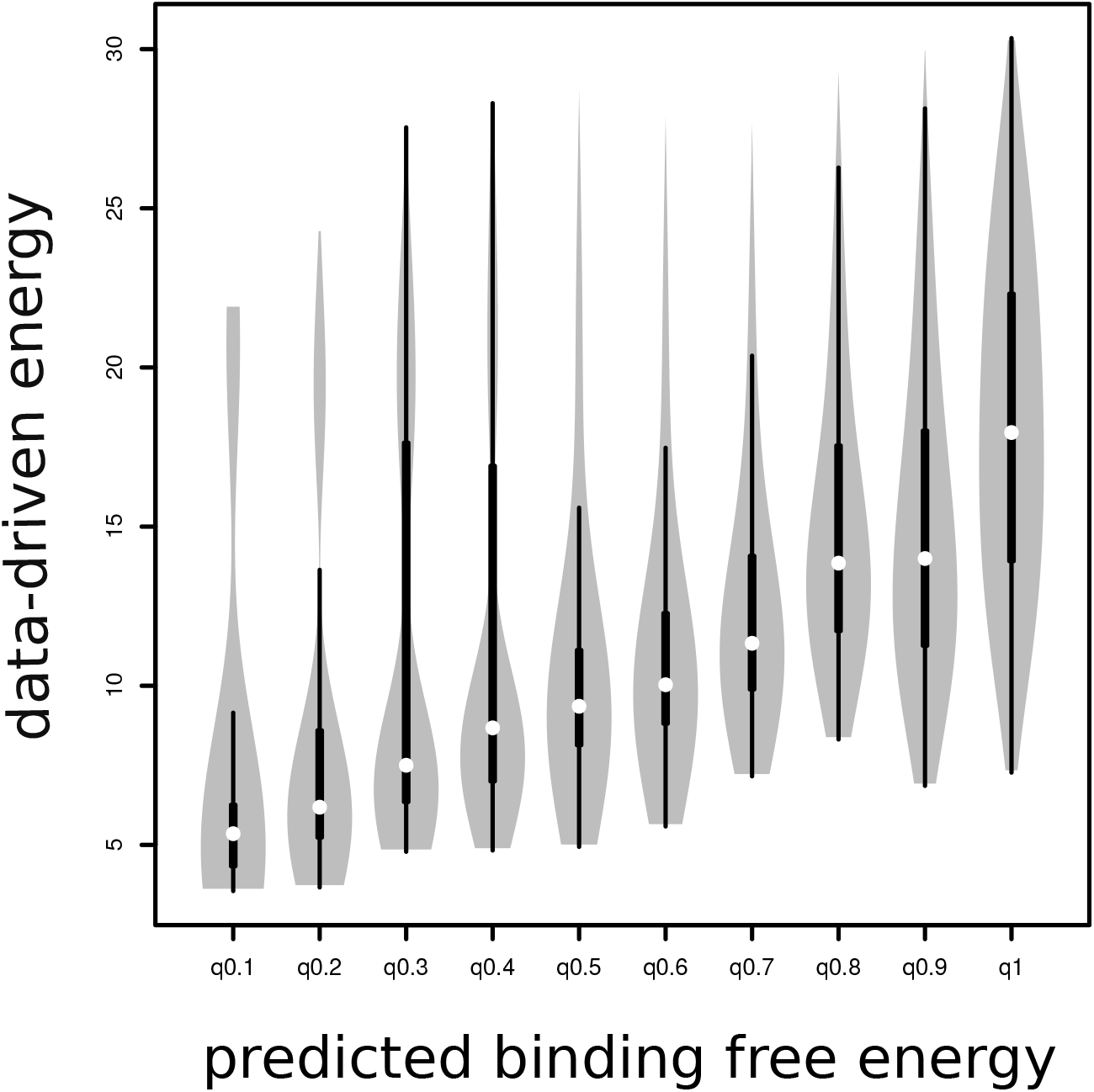
Data driven energy. Boxplots of data-driven energy values estimated for 502197 human 5’ss sequences for every decile of predicted dimerization energies against U1 smRNA. Dimerization free energy between the sequences of the donor sites and the complementary portion of the snRNA U1 was estimated using the RNAcofold program of the ViennaRNA 2.0 package [45], using default parameters.

**Sup Fig 6.**
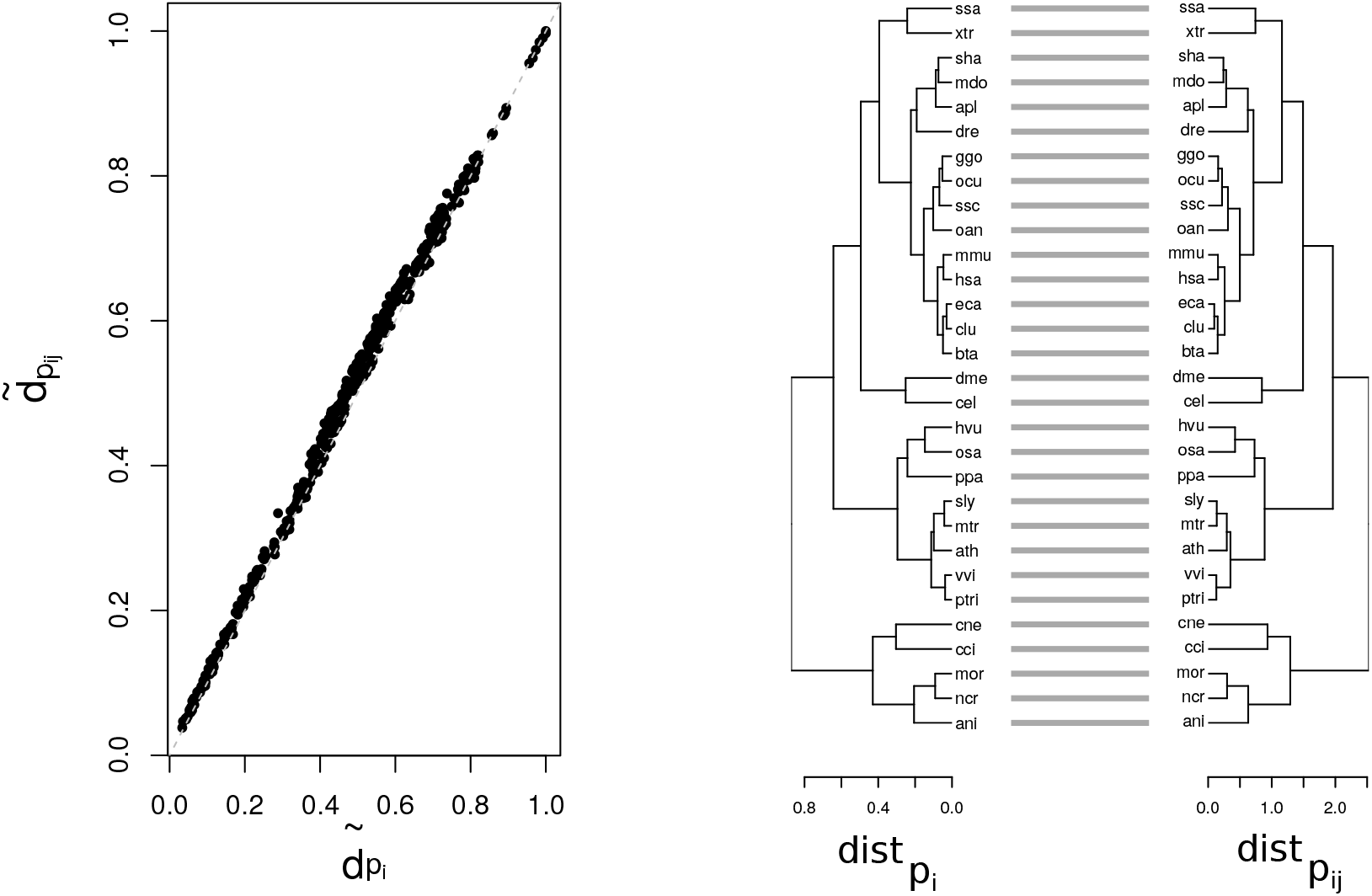
Distances inferred from one-site and two-site statistics. Left panel of this figure shows pairwise scaled euclidean distances between species estimated using *P*_*i*_ (x-axis) and *P*_*ij*_ (y-axis) information. The observed linear relationship and the perfectly matched tanglegram, showed at the right panel, highlight the strong correlation between these two metrics.

**Sup Fig 7.**
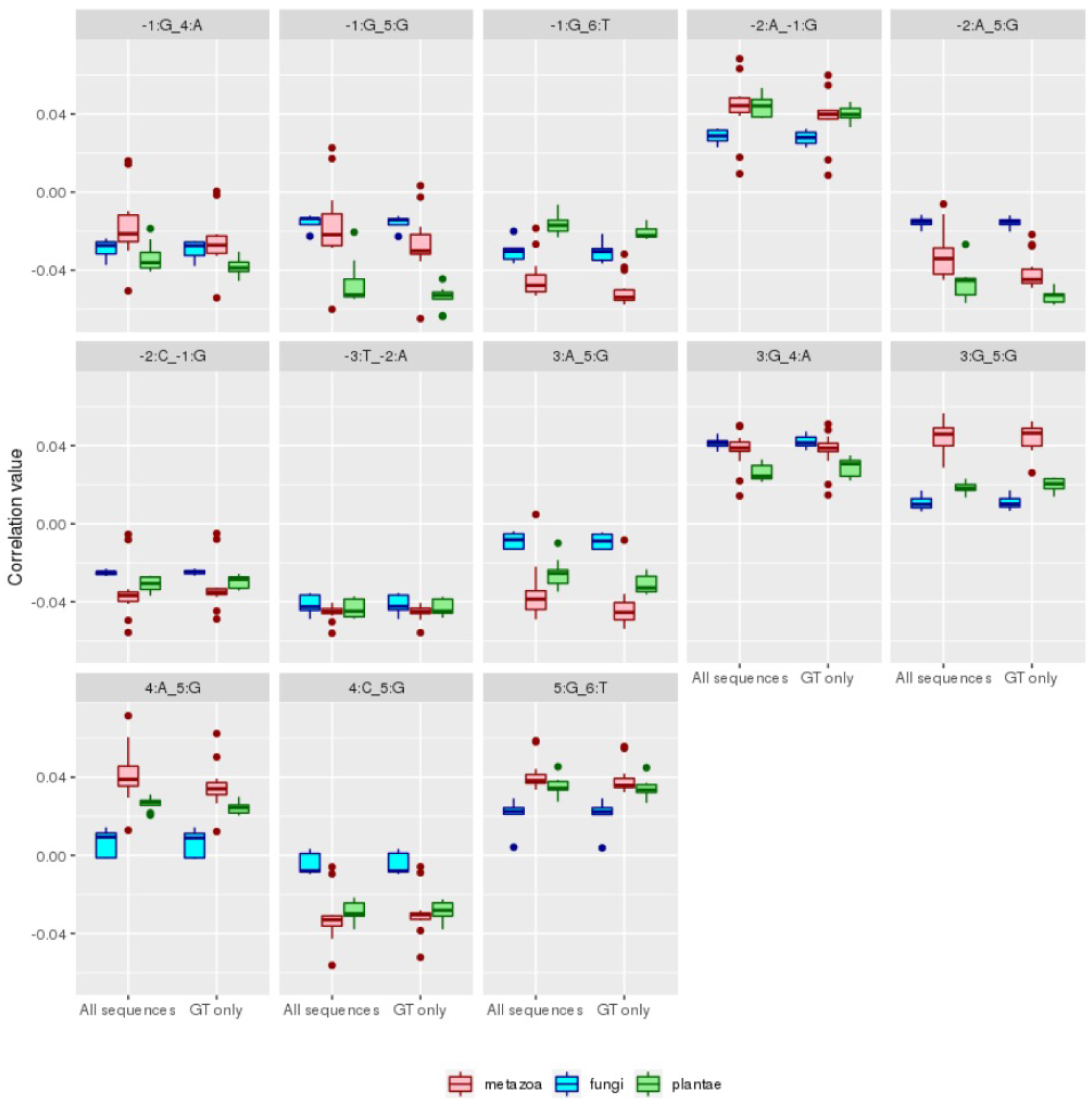
Di-nucleotide pairwise correlations. Each panel of the figure shows a set of boxplots summarizing the distribution of correlation values obtained for animal (red), fungi (blue) and plants (green) considering a specific di-nucleotide pair. The complete set of annotated 5’ splicing sequences, and just GT 5’ss were considered in the first and second boxplot triplets respectively.

**Sup Fig 8.**
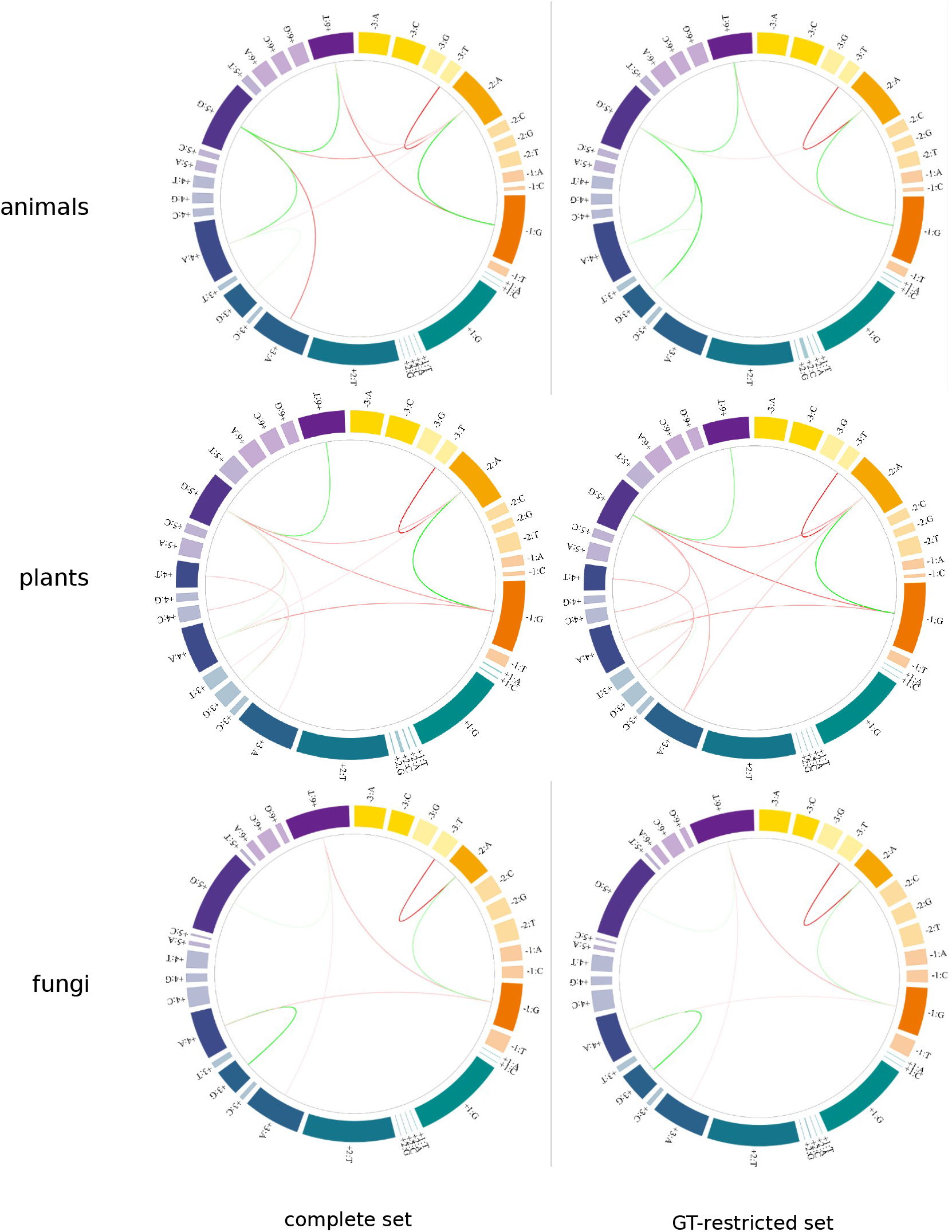
Circos. Circos diagrams for coupling patterns estimated for the complete set or GT-restricted donor sequences are displayed in the first and second column respectively. Results for animals, plants and fungi are shown in the first, second and third rows respectively.

**Sup Fig 9.**
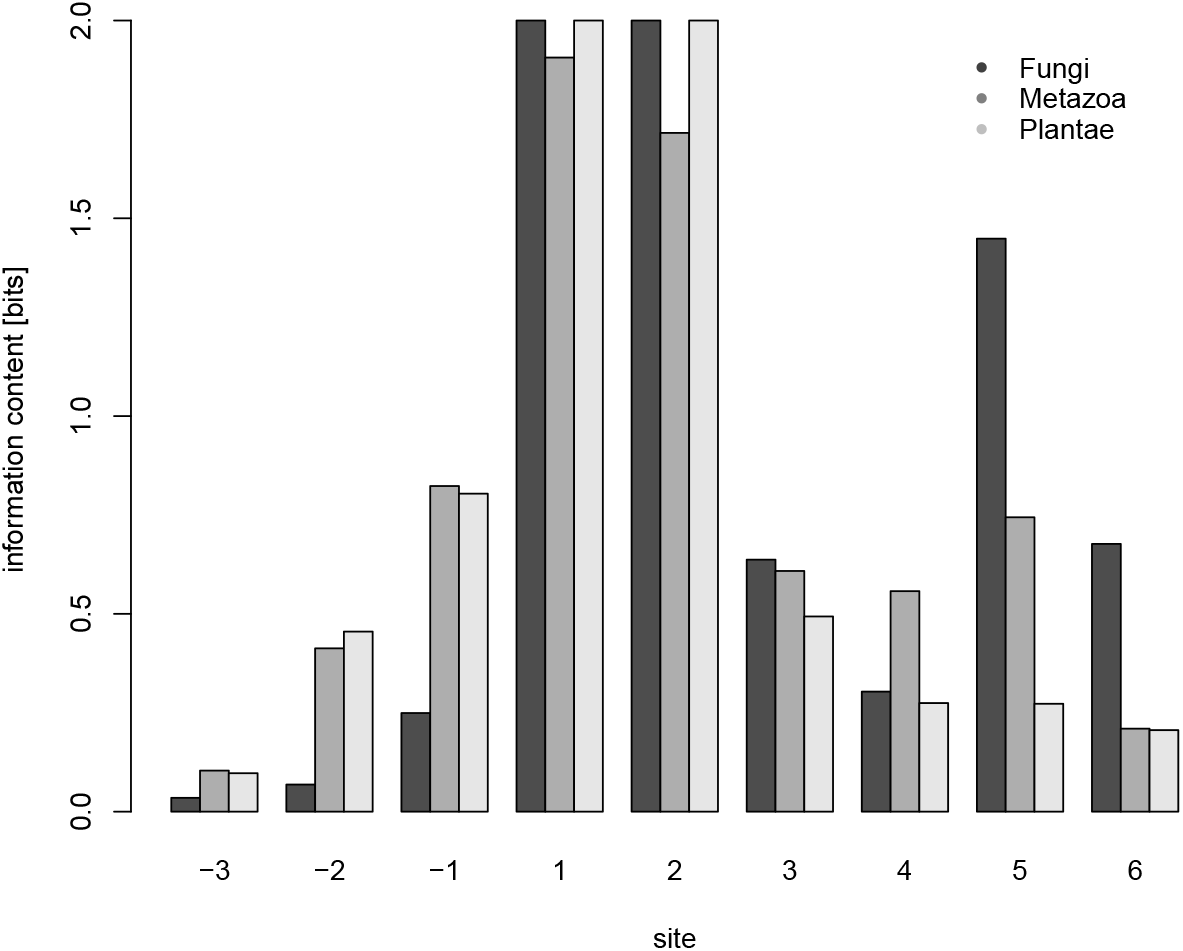
Information content. Barplot of information content per site for plants, animal and fungi.

**Supplementary Table 1.** Analyzed genomes. For each analyzed organism we reported, the assembly code, the total number of 5’ exon-intron boundaries, the number of different 5’ss sequences and the number of different 5’ss sequences presenting *GT* as the first two intronic bases.

| organism | code | assembly | 5'ss (unique) | 5'ss GT (unique) |
| --- | --- | --- | --- | --- |
| *Aspergillus nidulans* | ani | ASM1142v1 | 24563 (3954) | 24514 (3919) |
| *Coprinopsis cinerea okayama* | cci | CC3 | 61427 (4737) | 61325 (4670) |
| *Cryptococcus neoformans* | cne | cryp_neof.125.91.V1 | 36046 (2637) | 35220 (2435) |
| *Magnaporthe oryzae* | mor | MG8 | 22772 (3126) | 22567 (3039) |
| *Neurospora crassa* | ncr | NC12 | 16739 (2522) | 16533 (2423) |
| *Arabidopsis thaliana* | ath | TAIR10 | 136036 (11286) | 130069 (6803) |
| *Hordeum vulgare* | hvu | IBSCv2 | 499846 (48879) | 413160 (10107) |
| *Medicago truncatula* | mtr | MedtrA17.4.0 | 156335 (11396) | 148511 (6964) |
| *Oryza sativa* | osa | IRGSP-1.0 | 130702 (12688) | 124646 (8418) |
| *Physcomitrium patens* | ppa | Phypa V3 | 194684 (8310) | 191147 (6991) |
| *Populus trichocarpa* | ptri | Pop_tri.v3 | 179803 (10168) | 176766 (9285) |
| *Solanum lycopersicum* | sly | SL3.0 | 141263 (12562) | 135225 (8400) |
| *Vitis vinifera* | vvi | 12X | 108097 (9614) | 105266 (7713) |
| *Anas platythynchos* | apl | CAU_duck1.0 | 152272 (9075) | 146465 (5405) |
| *Bos taurus* | bta | ARS_UCD1.2 | 203086 (11919) | 194432 (6624) |
| *Canis lupus familiaris* | clu | CanFam3.1 | 215912 (12437) | 206340 (5521) |
| *Danio rerio* | dre | GRCz11 | 276776 (11723) | 268477 (6720) |
| *Equus caballus* | eca | EquCab3.0 | 216007 (10367) | 208399 (4984) |
| *Gorilla gorilla* | ggo | gorGor4 | 212738 (14620) | 200262 (7697) |
| *Homo sapiens* | hsa | GRChv38 | 502197 (12129) | 488939 (6206) |
| *Monodelphis domestica* | mdo | ASM229v1 | 208511 (14090) | 196940 (6970) |
| *Mus musculus* | mmu | GRChm39 | 391997 (9026) | 384141 (5369) |
| *Ornithorhynchus anatinus* | oan | mOrnAna1.p.v1 | 187119 (13517) | 175225 (6562) |
| *Oryctolagus cuniculus* | ocu | OryCun2.0 | 147142 (12204) | 138124 (5217) |
| *Sarcophilus harrisii* | sha | mSarHar1.11 | 227688 (10104) | 220325 (5637) |
| *Salmo salar* | ssa | ICSASGv2 | 516921 (28445) | 455502 (14209) |
| *Sus scrofa* | ssc | Sscrofa11.1 | 260642 (21094) | 243064 (8300) |
| *Xenopus tropicalis* | xtr | Xenopus_tropicalis_v9.1 | 231618 (20862) | 205430 (10794) |
| *Caenorhabditis elegans* | cel | WBcel235 | 127661 (7625) | 124619 (5351) |
| *Drosophila melanogaster* | dme | BDGP6.32 | 63121 (4023) | 62451 (3780) |

**Supplementary Table 2.** Conserved patterns (*γ* = 0.015) Mean interactions between different type of sites are shown for different organisms. EC, ENC, IC, and INC stand for exonic-consensus, exonic-non-consensus, intronic-consensus and intronic-non-consensus respectively.

| Species | IC-EC | IC-ENC | INC-EC | INC-ENC | IC-IC | IC-INC | INC-INC | EC-EC | ENC-EC | ENC-ENC |
| --- | --- | --- | --- | --- | --- | --- | --- | --- | --- | --- |
| cne | -0.08 | 0 | 0 | 0 | 0.34 | 0 | 0 | 0.92 | 0 | -0.17 |
| ani | -0.48 | 0 | 0 | 0 | 0.06 | 0 | 0 | 0.88 | 0 | 0 |
| ncr | -0.56 | 0 | 0 | 0 | 0.26 | 0 | 0 | 0.79 | 0 | 0 |
| mor | -0.29 | 0 | 0 | 0 | 0 | 0 | 0 | 0.96 | 0 | 0 |
| cci | -0.28 | 0 | 0 | 0 | 0.24 | 0.04 | 0 | 0.93 | 0 | 0 |
| ath | -0.37 | 0 | 0.02 | 0 | 0.19 | 0 | 0 | 0.91 | 0 | 0 |
| hvu | 0 | 0.13 | 0 | -0.08 | 0 | 0.94 | 0.19 | 0 | 0 | -0.23 |
| mtr | -0.56 | 0 | 0 | 0 | 0.07 | 0 | 0 | 0.82 | 0 | 0 |
| osa | -0.22 | 0 | 0 | 0 | 0.07 | 0.02 | -0.03 | 0.97 | 0 | 0 |
| ppa | -0.84 | 0 | 0 | 0 | 0.05 | 0 | -0.03 | 0.55 | 0 | 0 |
| ptri | -0.42 | 0 | 0 | 0 | 0.02 | 0 | 0 | 0.91 | 0 | 0 |
| sly | -0.33 | 0 | 0 | 0 | 0.23 | 0 | 0 | 0.92 | 0 | 0 |
| vvi | -0.34 | 0 | 0 | 0 | 0.01 | 0 | 0 | 0.94 | 0 | 0 |
| apl | -0.41 | 0 | 0 | 0 | 0.39 | 0 | 0 | 0.83 | 0 | 0 |
| bta | -0.27 | 0 | 0 | 0 | 0.59 | 0 | 0 | 0.76 | 0 | 0 |
| clu | -0.33 | 0 | 0 | 0 | 0.56 | 0 | 0 | 0.76 | 0 | 0 |
| dre | -0.34 | 0 | 0 | 0 | 0.24 | 0 | 0 | 0.91 | 0 | 0 |
| eca | -0.36 | 0 | 0 | 0 | 0.51 | 0 | 0 | 0.78 | 0 | 0 |
| ggo | -0.22 | 0 | 0 | 0 | 0.57 | 0 | 0 | 0.79 | 0 | 0 |
| hsa | -0.55 | 0 | 0 | 0 | 0.16 | 0 | 0 | 0.82 | 0 | 0 |
| mdo | -0.19 | 0 | 0 | 0 | 0.57 | 0 | 0 | 0.8 | 0 | 0 |
| mmu | -0.58 | 0 | 0 | 0 | 0.12 | 0 | 0 | 0.81 | 0 | 0 |
| oan | -0.23 | 0 | 0 | 0 | 0.61 | 0 | 0 | 0.76 | 0 | 0 |
| ocu | -0.27 | 0 | 0 | 0 | 0.59 | 0 | 0 | 0.76 | 0 | 0 |
| sha | -0.31 | 0 | 0 | 0 | 0.63 | 0 | 0 | 0.71 | 0 | 0 |
| ssa | 0.11 | 0 | -0.05 | 0 | 0.61 | 0 | 0 | 0.78 | 0 | 0 |
| ssc | -0.18 | 0 | 0 | 0 | 0.62 | 0 | 0 | 0.77 | 0 | 0 |
| xtr | 0.17 | 0 | -0.05 | 0 | 0.59 | 0 | 0 | 0.79 | 0 | 0 |
| dme | -0.97 | 0 | 0 | 0 | -0.26 | 0 | 0 | 0 | 0 | 0 |
| cel | -0.97 | 0 | 0.06 | 0 | -0.21 | 0 | 0 | 0 | 0 | -0.1 |

## Data availability

Donor sequence data, source code and in-house scripts can be found at https://github.com/chernolab/Mining5ss_SM.

## Acknowledgments

The authors would like to thank Rocio Espada, Ezequiel Petrillo, Anabella Srebrow and Hernan Dopazzo for helpful discussions.

## References

1. Nilsen TW. The spliceosome: The most complex macromolecular machine in the cell?; 2003. Available from: https://pubmed.ncbi.nlm.nih.gov/14635248/.

2. Dujardin G, Lafaille C, de la Mata M, Marasco LE, Muñoz MJ, Le Jossic-Corcos C, et al. How Slow RNA Polymerase II Elongation Favors Alternative Exon Skipping. Molecular Cell. 2014;54(4):683–690. doi:10.1016/j.molcel.2014.03.044.

3. Fong N, Kim H, Zhou Y, Ji X, Qiu J, Saldi T, et al. Pre-mRNA splicing is facilitated by an optimal RNA polymerase II elongation rate. Genes and Development. 2014;28(23):2663–2676. doi:10.1101/gad.252106.114.

4. Chen W, Moore MJ. Spliceosomes; 2015. Available from: http://www.cell.com/article/S096098221401553X/fulltexthttp://www.cell.com/article/S096098221401553X/abstracthttps://www.cell.com/current-biology/abstract/S0960-9822(14)01553-X.

5. Reddy ASN, Marquez Y, Kalyna M, Barta A, Reddy CW, N As, et al. Complexity of the Alternative Splicing Landscape in Plants. The Plant Cell. 2013;25:3657–3683. doi:10.1105/tpc.113.117523.

6. Rogozin IB, Carmel L, Csuros M, Koonin EV. Origin and evolution of spliceosomal introns. Biology Direct. 2012;7(1):1. doi:10.1186/1745-6150-7-11.

7. Daguenet E, Dujardin G, Valcárcel J. The pathogenicity of splicing defects: mechanistic insights into pre mRNA processing inform novel therapeutic approaches. EMBO reports. 2015;16(12):1640–1655. doi:10.15252/embr.201541116.

8. Kondo Y, Oubridge C, van Roon AMM, Nagai K. Crystal structure of human U1 snRNP, a small nuclear ribonucleoprotein particle, reveals the mechanism of 5<sup>I</sup> splice site recognition. eLife. 2015;4(e04986):805–817. doi:10.7554/eLife.04986.

9. Ast G. How did alternative splicing evolve? Nature Reviews Genetics. 2004;5(10):773–782. doi:10.1038/nrg1451.

10. Roca X, Krainer AR, Eperon IC. Pick one, but be quick: 5<sup>I</sup> splice sites and the problems of too many choices. Genes and Development. 2013;27(2):129–144. doi:10.1101/gad.209759.112.

11. Artemyeva-Isman OV, Porter ACG. U5 snRNA Interactions With Exons Ensure Splicing Precision. Frontiers in Genetics. 2021;12(July):1–33. doi:10.3389/fgene.2021.676971.

12. Wong MS, Kinney JB, Krainer AR. Quantitative Activity Profile and Context Dependence of All Human 5<sup>I</sup> Splice Sites. Molecular Cell. 2018;71(6):1012–1026.e3. doi:10.1016/J.MOLCEL.2018.07.033/ATTACHMENT/98398431-5D1A-41EC-B760-68671B576977/MMC1.PDF.

13. Stephens RM, Schneider TD. Features of spliceosome evolution and function inferred from an analysis of the information at human splice sites. Journal of Molecular Biology. 1992;228(4):1124–1136. doi:10.1016/0022-2836(92)90320-J.

14. Sverdlov AV, Rogozin IB, Babenko VN, Koonin EV. Evidence of Splice Signal Migration from Exon to Intron during Intron Evolution. Current Biology. 2003;13:2170–2174. doi:10.1016/j.cub.2003.12.003.

15. Iwata H, Gotoh O. Comparative analysis of information contents relevant to recognition of introns in many species. BMC Genomics. 2011;12(1):45. doi:10.1186/1471-2164-12-45.

16. Thanaraj TA, Robinson AJ. Prediction of exact boundaries of exons. Briefings in bioinformatics. 2000;1(4):343–356. doi:10.1093/bib/1.4.343.

17. Carmel I, Tal S, Vig I, Ast G. Comparative analysis detects dependencies among the 5<sup>I</sup> splice-site positions. Rna. 2004;10(5):828–840. doi:10.1261/rna.5196404.

18. Sahashi K, Masuda A, Matsuura T, Shinmi J, Zhang Z, Takeshima Y, et al. In vitro and in silico analysis reveals an efficient algorithm to predict the splicing consequences of mutations at the 5??? splice sites. Nucleic Acids Research. 2007;35(18):5995–6003. doi:10.1093/nar/gkm647.

19. Denisov S, Bazykin G, Favorov A, Mironov A, Gelfand M. Correlated evolution of nucleotide positions within splice sites in mammals. PLoS ONE. 2015;10(12):1–24. doi:10.1371/journal.pone.0144388.

20. Yeo G, Hoon S, Venkatesh B, Burge CB. Variation in sequence and organization of splicing regulatory elements in vertebrate genes. Proceedings of the National Academy of Sciences. 2004;101(44):15700–15705. doi:10.1073/pnas.0404901101.

21. Bialek W, Ranganathan R. Rediscovering the power of pairwise interactions. arXiv preprint. 2007;(q-bio.QM):1–8.

22. De Martino A, De Martino D. An introduction to the maximum entropy approach and its application to inference problems in biology; 2018.

23. Roudi Y, Aurell E, Hertz Ja. Statistical physics of pairwise probability models. Front Comput Neurosci. 2009;3(November):22. doi:10.3389/neuro.10.022.2009.

24. Figliuzzi M, Barrat-Charlaix P, Weigt M. How pairwise coevolutionary models capture the collective residue variability in proteins? Molecular Biology and Evolution. 2018;35(4):1018–1027. doi:10.1093/molbev/msy007.

25. Bialek W, Cavagna A, Giardina I, Mora T, Silvestri E, Viale M, et al. Statistical mechanics for natural flocks of birds. Proceedings of the National Academy of Sciences of the United States of America. 2012;109(13):4786–4791. doi:10.1073/pnas.1118633109.

26. Espada R, Parra RG, Mora T, Walczak AM, Ferreiro DU. Inferring repeat-protein energetics from evolutionary information. PLoS Computational Biology. 2017;13(6):1–16. doi:10.1371/journal.pcbi.1005584.

27. R Jun Base: a database of RNA splice junctions in human normal and cancerous tissues Qin. Nucleic Acids Research. 2021;49(D1):D201––D211. doi:10.1093/nar/gkaa1056.

28. Maddison WP, Montgomery S. Null models for the number of evolutionary steps in a character on a phylogenetic tree. Evolution. 1991;(45):1184–1197.

29. D S. Minimal Mutation Trees of Sequences. Journal Applied Math. 1975;(28(1)):35–42.

30. P Sk. phangorn: phylogenetic analysis in R. Bioinformatics. 2011;27(4):592–593.

31. Galil T. dendextend: an R package for visualizing, adjusting, and comparing trees of hierarchical clustering. Bioinformatics. 2015;doi:10.1093/bioinformatics/btv428.

32. Fowlkes CL E B; Mallows. A Method for Comparing Two Hierarchical Clusterings. Journal of the American Statistical Association. 1983;(78(383)):553–42.

33. Moyer DC, Larue GE, Hershberger CE, Roy SW, Padgett RA. Comprehensive database and evolutionary dynamics of U12-type introns. Nucleic Acids Research. 2020;48(13):7066–7078. doi:10.1093/nar/gkaa464.

34. Schwartz SH, Silva J, Burstein D, Pupko T, Eyras E, Ast G. Large-scale comparative analysis of splicing signals and their corresponding splicing factors in eukaryotes. Genome Research. 2008;18(1):88–103. doi:10.1101/gr.6818908.

35. Joseph F. Inferring Phylogenies. Sinauer Associates; 2004.

36. Chaudhary S, Khokhar W, Jabre I, Reddy ASN, Byrne LJ, Wilson CM, et al. Alternative splicing and protein diversity: Plants versus animals. Frontiers in Plant Science. 2019;10(June):1–14. doi:10.3389/fpls.2019.00708.

37. Wang ET, Sandberg R, Luo S, Khrebtukova I, Zhang L, Mayr C, et al. Alternative isoform regulation in human tissue transcriptomes. Nature. 2008;456(7221):470–476. doi:10.1038/nature07509.

38. Chamala S, Feng G, Chavarro C, Barbazuk WB. Genome-wide identification of evolutionarily conserved alternative splicing events in flowering plants. Frontiers in Bioengineering and Biotechnology. 2015;3(MAR):33. doi:10.3389/fbioe.2015.00033.

39. Pan Q, Shai O, Lee LJ, Frey BJ, Blencowe BJ. Deep surveying of alternative splicing complexity in the human transcriptome by high-throughput sequencing. Nature Genetics. 2008;40(12):1413–1415. doi:10.1038/ng.259.

40. Merkin J, Russell C, Chen P, Burge CB. Evolutionary dynamics of gene and isoform regulation in mammalian tissues. Science. 2012;338(6114):1593–1599. doi:10.1126/science.1228186.

41. Vosseberg J, Snel B. Domestication of self-splicing introns during eukaryogenesis: the rise of the complex spliceosomal machinery. Biology Direct. 2017;doi:10.1186/s13062-017-0201-6.

42. Carvalho RF, Szakonyi D, Simpson CG, Barbosa ICR, Brown JWS, Baena-González E, et al. The arabidopsis SR45 splicing factor, a negative regulator of sugar signaling, modulates SNF1-related protein kinase 1 stability. Plant Cell. 2016;28(8):1910–1925. doi:10.1105/tpc.16.00301.

43. Laloum T, Martín G, Duque P. Alternative Splicing Control of Abiotic Stress Responses. Trends in Plant Science. 2018;23(2):140–150. doi:10.1016/j.tplants.2017.09.019.

44. Jia J, Long Y, Zhang H, Li Z, Liu Z, Zhao Y, et al. Post-transcriptional splicing of nascent RNA contributes to widespread intron retention in plants. Nature Plants. 2020;6(7):780–788. doi:10.1038/s41477-020-0688-1.

45. Lorenz R, Bernhart SH, Höner Zu Siederdissen C, Tafer H, Flamm C, Stadler PF, et al. ViennaRNA Package 2.0; 2011. Available from: http://www.tbi.univie.ac.at/RNA.

